# Novel two-stage processes for optimal chemical production in microbes

**DOI:** 10.1101/803023

**Authors:** Kaushik Raj, Naveen Venayak, Radhakrishnan Mahadevan

## Abstract

Microbial metabolism can be harnessed to produce a broad range of industrially important chemicals. Often, three key process variables: **T**iter, **R**ate and **Y**ield (**TRY**) are the target of metabolic engineering efforts to improve microbial hosts toward industrial production. Previous research into improving the TRY metrics have examined the efficacy of having distinct growth and production stages to achieve enhanced productivity. However, these studies assumed a switch from a maximum growth to a maximum production phenotype. Hence, phenotypes with intermediate growth and chemical production for the growth and production stages of two-stage processes are yet to be explored. The impact of reduced growth rates on substrate uptake adds to the need for intelligent choice of operating points while designing two-stage processes. In this work, we develop a computational framework that scans the phenotypic space of microbial metabolism to identify ideal growth and production phenotypic targets, to achieve optimal TRY targets. Using this framework, with *Escherichia coli* as a model organism, we compare two-stage processes that use dynamic pathway regulation, with one-stage processes that use static intervention strategies, for different bioprocess objectives. Our results indicate that two-stage processes with intermediate growth during the production stage always result in optimal TRY values even in cases where substrate uptake is limited due to reduced growth during chemical production. By analyzing the flux distributions for the production enhancing strategies, we identify key reactions and reaction subsystems that require perturbation to achieve a production phenotype for a wide range of metabolites in *E. coli*. Interestingly, flux perturbations that increase phosphoenolpyruvate and NADPH availability are enriched among these production phenotypes. Furthermore, reactions in the pentose phosphate pathway emerge as key control nodes that function together to increase the availability of precursors to most products in *E. coli*. The inherently modular nature of microbial metabolism results in common reactions and reaction subsystems that need to be regulated to modify microbes from their target of growth to the production of a diverse range of metabolites. Due to the presence of these common patterns in the flux perturbations, we propose the possibility of a universal production strain.

## 1 Introduction

The use of microbes for the production of chemicals through metabolic engineering has garnered significant interest in the past few decades. The naturally modular arrangement of metabolic networks makes microbes amenable for use as chemical production platforms^1^. Metabolic networks have a bow-tie architecture which allows a large number of metabolites to be produced from a few universal precursors^2^. This has allowed us to successfully engineer microbes to be biocatalysts for the production of a wide range of commodity chemicals^3,4^, pharmaceuticals^5,6^, biofuels,^7,8^ and other natural and non-natural compounds^9^. While some such production processes have been successful at an industrial scale^10,11^, large strain development costs and scale-up issues could deem many processes economically infeasible^12,13^. Given the cost of a target feedstock and product, the feasibility of industrial fermentation processes is typically determined by three process metrics - **T**iter: concentration of product at the end of a fermentation batch (given in *mmol/L* or *g/L* of product), **R**ate/productivity: the rate of product secretion (given in *mmol/L.h* or *g/L.h* of product), and **Y**ield: the amount of product produced per unit amount of substrate (given in *mmol* product*/mmol* substrate or *g* product*/g* substrate) - collectively termed the **TRY** metrics^14^. Titer and yield affect the operating expenditure of the process by impacting product separation and substrate costs respectively, while productivity affects the capital expenditure by determining the scale of the reactor required. Microbial production processes undergo several rounds of strain, pathway and process optimization to reach acceptable TRY targets^15,16^.

Recent advancements in genome scale reconstructions of microbial metabolism have augmented the process of strain development. Wild-type microbial strains have typically evolved to grow at maximal rates, directing little carbon flux towards production of target compounds^17^. Metabolic engineering attempts to alter the phenotype (or operating point in a production envelope) of these strains to enhance target chemical production by throttling flux through growth associated reactions and/or tuning native metabolism to balance pathway energy and cofactor requirements. Given a stoichiometric model of microbial metabolism and substrate/nutrient uptake rates, the feasible range of chemical production characteristics in a microbial strain can be visualized using its production envelope and yield envelope^18^, which map the maximum product flux and maximum product yield respectively, at all possible growth rates of the microbe (Figure 1). Strain engineering strategies to improve TRY metrics can be broadly classified into static and dynamic pathway engineering strategies. Static pathway engineering involves making gene deletions that either couple the production of a target compound with the microorganism’s growth^19^ or, simply redirect more carbon flux through production pathways. These strategies are typically implemented as one-stage (OS) production processes where the strain remains at a single operating point throughout the course of the process (Figure 1a). Such processes result in higher yield by ensuring high relative pathway flux. Recently, there has been an increased interest in dynamic pathway engineering, which involves temporally controlling carbon flux through growth and production pathways. This can be achieved through the use of biological logic or sensor and actuator systems composed of cellular components^20–22^. These strategies are implemented as two-stage (TS) production processes which start with cells in their growth stage and at some point during the fermentation, production pathway genes are expressed to switch to the production stage (Figure 1b). Such a decoupling of growth and production stages is thought to reduce batch times by reaching maximal biomass concentrations faster and thereby increase productivity^23,24^.

**Figure 1:**
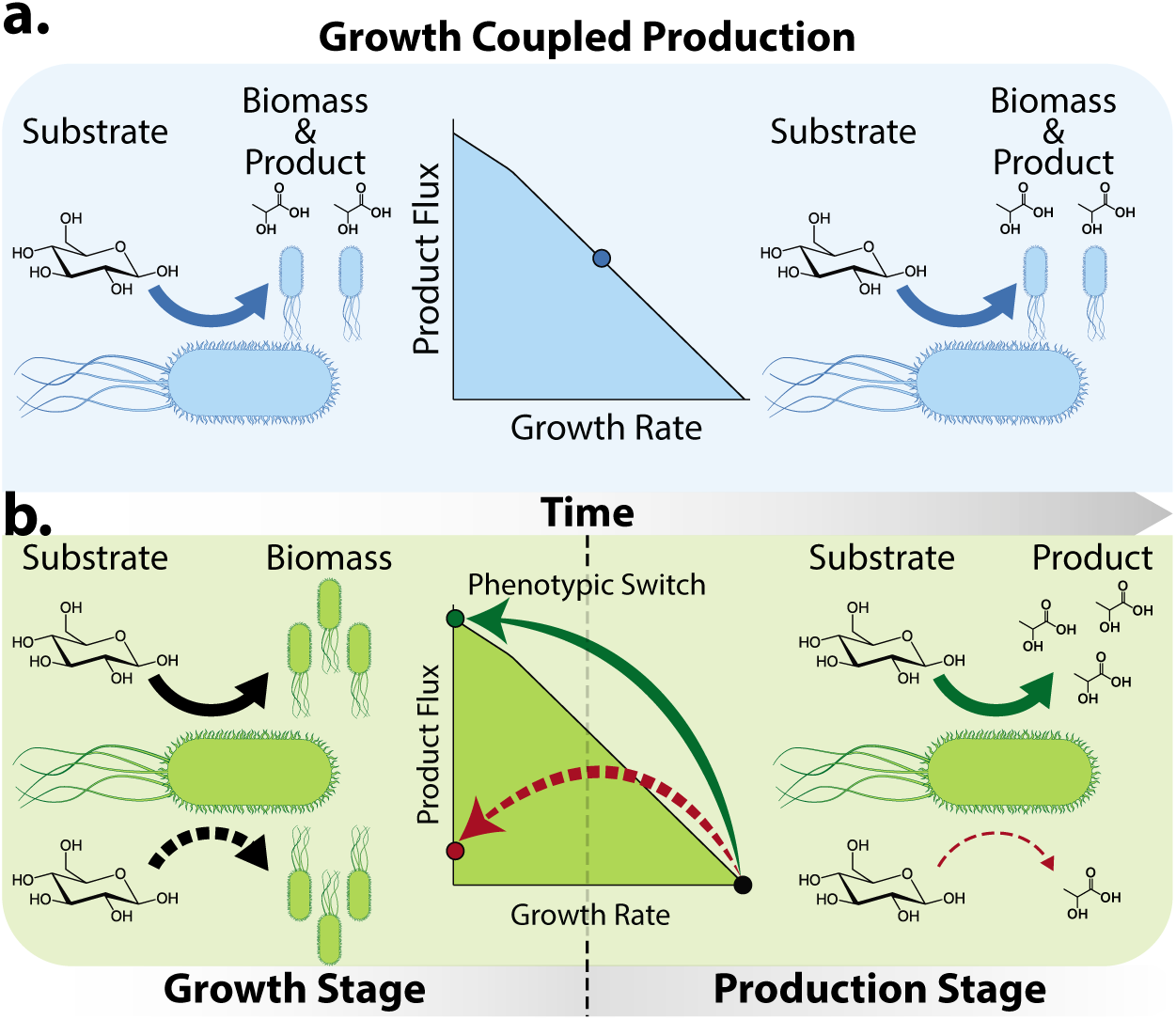
Static vs dynamic pathway engineering. Strain engineering strategies can be classified into **a.** Static engineering strategies where genetic perturbations that allow cells to grow and produce the target compound simultaneously (growth coupled production) are implemented. This enables the cells to produce the compound in a one-stage process. **b.** Dynamic engineering strategies where growth and production pathways are decoupled temporally. In such strategies, the process starts with a growth stage, accumulating biomass and switches over to a production stage to produce the target compound. Reduced substrate uptake during the production stage can result in lower product flux (dotted lines) than that expected assuming constant substrate uptake rates (solid lines). Hypothetical operating points for each production strategy are shown in the respective production envelopes.

While stoichiometric models are effective for determining relative production metrics such as yield, absolute metrics such as end-titer and productivity are also governed by variations in substrate uptake rates. Metabolic models with constant substrate uptake rates are routinely used for simulations to monitor metabolite production rates at different phases of metabolism. The impact of reduced substrate uptake rate during stationary phase metabolism^25^ is often overlooked while designing microbial production processes. Studies have shown that the rate of glucose uptake varies significantly depending on the genotype of the strain and phase of metabolism^25–31^ (Supplementary Figure S1a). A drop in substrate uptake rate during the production stage of a TS process would result in reduced product flux and therefore lower productivity, defeating the purpose behind designing such a process (dotted lines in Figure 1b). This effect was recently shown in a theoretical study that compared the performance of TS and OS processes for D-lactic acid production in *E. coli*^32^. This study showed that reduced substrate uptake rates can limit the advantages of a TS process and in some cases, there is a very narrow range of conditions where a TS process can outperform an OS process. One way to resolve this issue is to employ techniques to increase stationary phase substrate uptake, such as engineering ATP futile cycles to expand the range of conditions in which TS processes offer enhanced productivity. This study and many others consider only TS processes that switch from wild-type growth to a non-growing production phenotype during the stationary phase of metabolism. However, given the interplay between substrate uptake and growth rates, phenotypes with intermediate growth and production could hold significant value. Intermediate phenotypes have been examined in the past to identify operating points that result in balanced TRY values in OS processes^33^.

In this work, we compare TS and OS production processes that make use of the entire range of feasible production operating points rather than those with maximum growth or maximum product yield to identify better production phenotypes for bioprocesses. To this end, we develop **mcPECASO** (**m**icrobial **c**hemical **P**roduction **E**nhancement via **C**omplete **A**nalysis of **S**witchable **O**perating-points) - a modular computational framework that can compare microbial production processes. mcPECASO uses a two-stage dynamic flux balance analysis to determine the process metrics obtainable using hypothetical operating points calculated within the solution space of a microbe’s metabolic model. With this information, the framework can determine the best process type and optimal phenotypic choices that result in the maximum value of a predetermined objective. We use mcPECASO to discover enhanced TS processes that result in high TRY values while considering the substrate uptake effects of reduced growth. We also identify flux perturbations that occur consistently in production strategies for all natural products, giving rise to the possibility of a universal chassis for metabolic engineering.

## 2 Materials & Methods

The methods used in the mcPECASO formulation and the analysis of production stage fluxes are outlined below.

### 2.1 mcPECASO formulation

The mcPECASO workflow formulated in this study is briefly summarized in Figure 2. The overall goal in mcPECASO was to identify optimal phenotypic combinations for TS chemical production processes and compare TS processes with OS processes towards a user defined bioprocess objective for a broad range of host strains and target compounds. Therefore, we formulated mcPECASO as a modular framework where the choice of host organism, target compound and fermentation parameters can be readily modified to suit the user’s requirements. The individual steps involved in this formulation are described below.

**Figure 2:**
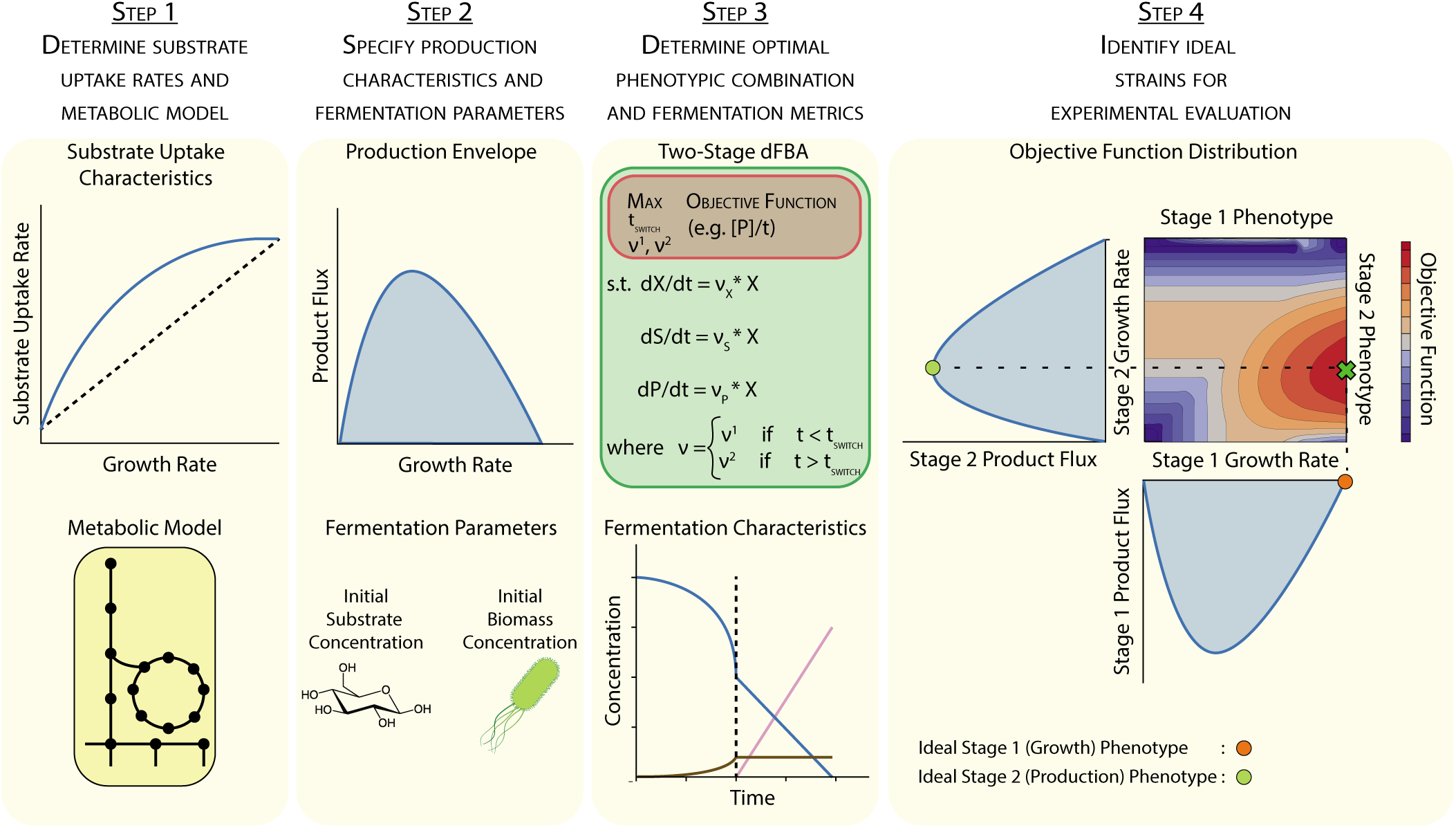
mcPECASO workflow. **Step 1** - A metabolic reconstruction of the microorganism and an approximation of the substrate uptake characteristics at different growth rates are given as inputs to the formulation. Currently, mcPECASO accepts all COBRA compatible metabolic models and any mathematical function to model substrate uptake variations. **Step 2** - A realistic production envelope is generated for the product of interest. This entails maximizing the product flux at all possible growth rates of the organism while considering reduced substrate uptake limitation at low growth rates. Fermentation start parameters such as initial biomass and substrate concentrations are given as inputs. **Step 3** - A two-stage dynamic flux balance analysis that maximizes a user-defined objective function (any combination of the process metrics - productivity, yield, and titer) is conducted to determine the process metrics, and optimal combination of phenotypes. **Step 4** - A distribution of process metrics for all two-stage processes is plotted and strains for experimental evaluation are chosen. The optimal phenotype for each stage has been projected onto the respective production envelope.

#### 2.1.1 Establishing substrate uptake characteristics

The first step towards analyzing the performance of chemical production strategies using a metabolic model is establishing a relationship between the growth rate and the substrate uptake rate in the organism being studied. There are several studies that have attempted to elucidate the relationship between substrate uptake and growth rates in *E. coli* - the host of our choice. Many of these have examined this relationship using a chemostat under glucose limiting conditions^34–37^. Under these conditions, the rate of glucose uptake is limited by the dilution rate prevailing in the reactor and not by the effects of genetic perturbations in the cells. Therefore, for our analysis we only considered studies with batch fermentations under glucose excess conditions to identify the relationship between growth and substrate uptake^25–31^ (Data attached as supplementary file). Furthermore, we restricted our analysis to the MG1655 strain of *E. coli* due to its ubiquitous use in research and industrial biotechnology (Supplementary Figure S1b).

Preliminary analysis of growth dependent substrate uptake rates in *E. coli* MG1655 did not reveal a one-to-one relationship. This is possibly due to the fact that the data-points collected (Supplementary Figure S1b) belong to strains with different gene deletions and each deletion potentially affects substrate uptake and growth rates through a different mechanism. Glucose uptake regulation in *E. coli* is quite complex and controlled by global transcriptional regulators and more directly by intracellular metabolites through allosteric regulation^38^. Direct allosteric regulation of glucose uptake has been shown to result from a build-up of intracellular metabolites involved in the citric acid cycle such as *α*-ketoglutarate, which competitively inhibits one of the enzymes involved in the phosphotransferase system^39–41^. With the assumption that the levels of intracellular metabolites increase transiently upon a switch to lower growth rates due to carbon flux being partitioned towards production pathways, and that the extent of this increase depends on the extent of growth rate reduction, we propose that glucose uptake rates likely obey a saturation type model where glucose uptake rate increases with growth rates and saturates at its maximum value at wild-type growth. Therefore, we chose to use a logistic curve - the most commonly used saturation type model as an approximation of substrate uptake variation with growth (shown in Supplementary Text 1.1). The parameter values for this curve were chosen such that the model would match experimentally observed substrate uptake rates at wild-type growth and stationary phase growth, while resulting in the least possible substrate uptake rates at all intermediate growth rates, to represent a worst case scenario for substrate uptake inhibition. Higher values of substrate uptake at intermediate growth rates will not affect the conclusions of this study, as will be seen later.

#### 2.1.2 Determining a realistic production envelope

The next step is to determine the range of feasible product secretion rates for the strain over its entire growth range. In this study, we refer to a metabolic mode of an organism, represented by a unique combination of the possible growth and product secretion rates within the solution space of its metabolic model, as an operating point or ‘phenotype’. At intermediate growth rates, between the minimum and maximum values for the organism, determined from the metabolic model, we derive the corresponding substrate uptake rates using the relationship established in the previous step. Then, at each value of growth rate, we calculate the feasible range of product secretion rate or the production envelope, by constraining the growth and substrate uptake reactions to the required values and maximizing the secretion rate of the metabolite of interest. This is in contrast to traditional production envelopes where substrate uptake rates are assumed to be constant. Hence, while traditional production envelopes result in maximum product secretion at zero growth, the substrate uptake limited production envelope results in maximum product secretion at some intermediate value of growth rate. We believe that this novel mapping of product secretion to growth rates is a more realistic simulation of actual production rates that can be expected in mutant strains.

#### 2.1.3 Two-stage dynamic flux balance analysis

Dynamic flux balance analysis (dFBA)^42^ can be used to obtain process metrics for a fermentation process by simulating substrate, biomass and product concentrations over the course of a fermentation batch, provided that initial concentrations of these species are known. This is done by using ordinary differential equations to simulate changes in the concentration of relevant species using their fluxes obtained from a metabolic model. Here, we modify the dFBA formulation to allow for a phenotypic switch between the two stages of a TS fermentation process by using distinct biomass, substrate and product fluxes in the two stages as seen in Eq. 1d. In this equation, [*X*], [*S*], and [*P*] represent the biomass, substrate, and product concentrations respectively and *v*_*n,X*_, *v*_*n,S*_, and *v*_*n,P*_ are their corresponding production fluxes in the metabolic model for each stage. The biomass production flux (*v*_*n,X*_) is set by the outer optimization problem (Eq. 1a), which is described in the following section. Substrate flux (*v*_*n,S*_) corresponding to this value of *v*_*n,X*_ is calculated using the relationship established previously. The maximum possible product flux (*v*_*n,P*_) given these constraints is obtained in an inner optimization problem by performing flux balance analysis on a metabolic model by setting the product flux to be the objective and constraining the biomass and substrate fluxes to required values (Eq. 1e).

#### 2.1.4 Optimizing bioprocess objective

The goal of the optimization framework is to maximize either one or a weighted combination of the fermentation metrics - productivity, yield and titer (Eq. 1a), which can be readily calculated from the substrate and product concentrations at the end fo the fermentation batch obtained from the dynamic flux balance analysis formulation shown in Eq. 1d. This is achieved by varying the phenotypes (represented by the fluxes - *v*_1_ and *v*_2_) for the two stages, such that these phenotypes lie on the production envelope. More specifically, the biomass fluxes for each stage - *v*_1,*X*_ and *v*_2,*X*_ are varied in the outer optimization problem and the corresponding substrate and product fluxes are calculated as described previously. An additional design variable that affects the outcome of the fermentation batch is the time of switching between the two stages (*t*_*switch*_), which can vary between zero and a user-defined maximum value. In addition to the constraints on the design variables (Eq. 1b), we added optional constraints on the fermentation metrics which can be used to specify a minimum required productivity/yield/titer for the optimization problem (Eq. 1c). The reasons for choosing these objectives will be elaborated in the Results & Discussion section.

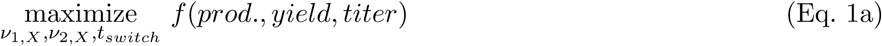

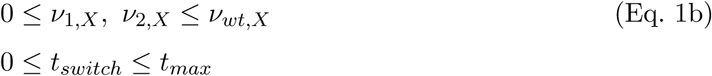

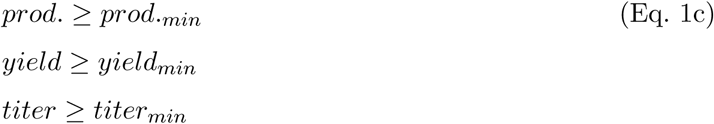

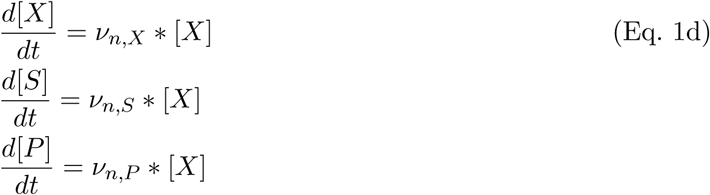

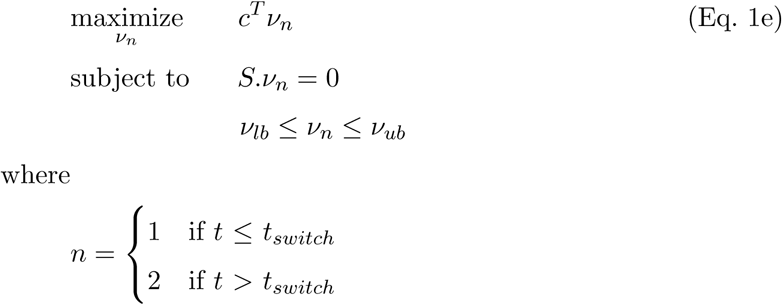

#### 2.1.5 Packaging and availability

The mcPECASO framework is written as a python package that accepts COBRApy^43^ compatible metabolic models. The modular nature of this package allows users to select a metabolic model, fermentation start parameters and, substrate uptake characteristics with ease. In order to reduce run times, we allowed for the optimization and dFBA calculations shown in Eq. 1 to be run parallelly on multi-core and multi-processor systems. The optimization problem was implemented using the COBYLA method in the optimization package of SciPy library in Python. The mcPECASO framework can be installed and run on any system with a working Python 3 distribution. The framework, along with installation instructions are available on GitHub^44^.

### 2.2 Implementation

All analyses were conducted using the COBRApy^43^ and Cameo^45^ packages on a Python 3.7 distribution. *E. coli* ‘s genome scale metabolic reconstruction - iJO1366^46^ was used to perform all simulations to compare the two fermentation strategies. Unless otherwise specified, fermentation batches were started with 500 mM (*≈* 90 *g/L*) of D-glucose as the substrate and 0.05 g/L of biomass. These values are in the range of required substrate and biomass concentrations to achieve acceptable TRY targets^15^. Three different bioprocess objectives were used to compare TS processes to OS processes in *E. coli* :

- *Objective I* : maximize productivity
- *Objective II* : maximize productivity with yield above a certain threshold (75% of the maximum yield unless otherwise specified)
- *Objective III* : maximize yield with productivity above 2 *g/L.h*

The reasons for choosing these objectives will be elaborated in the Results & Discussion section. Wildtype flux distributions were obtained through parsimonious flux balance analysis (pFBA)^47^. In order to minimize differences among the production stage flux distributions for all the products analyzed, we used the inherent redundant nature of metabolic networks to limit the number of different reactions involved in these flux distributions. This was done through several rounds of flux variability analysis (FVA)^48^ where reactions that can be constrained to zero for all products were successively removed. In the end, we obtained flux distributions for each product that only involved reactions that are either absolutely necessary for the production of that compound or are used im all or many of the other products analyzed. FVA and pFBA were run on the metabolic model using the IBM ILOG CPLEX (v12.9) solver and data visualization was performed using the plotly package. The source code used to perform the various analyses and generate figures used in this article are available on a GitHub repository^49^.

## 3 Results & Discussion

### 3.1 Case Study: Production of D-Lactic Acid in *E. coli*

We applied the newly formulated mcPECASO framework to predict strategies that maximize the productivity (*Objective I* described in the Materials & Methods section) of D-lactic acid production in *Escherichia coli*, starting with 500 *mM* of glucose as substrate and 0.05 *g/L* biomass. First, we examined the process under constant substrate uptake conditions i.e. assuming that the substrate uptake is unaffected by growth and other metabolic perturbations (Figure 3a). In these conditions, since the product flux and yield are highest when there is no growth, the best TS process for productivity is the traditional TS process, where the strain is allowed to switch from a maximum growth (wild-type) to a zero-growth phenotype (Figure 3b). As expected, the traditional TS process has a much higher productivity than the best OS process.

**Figure 3:**
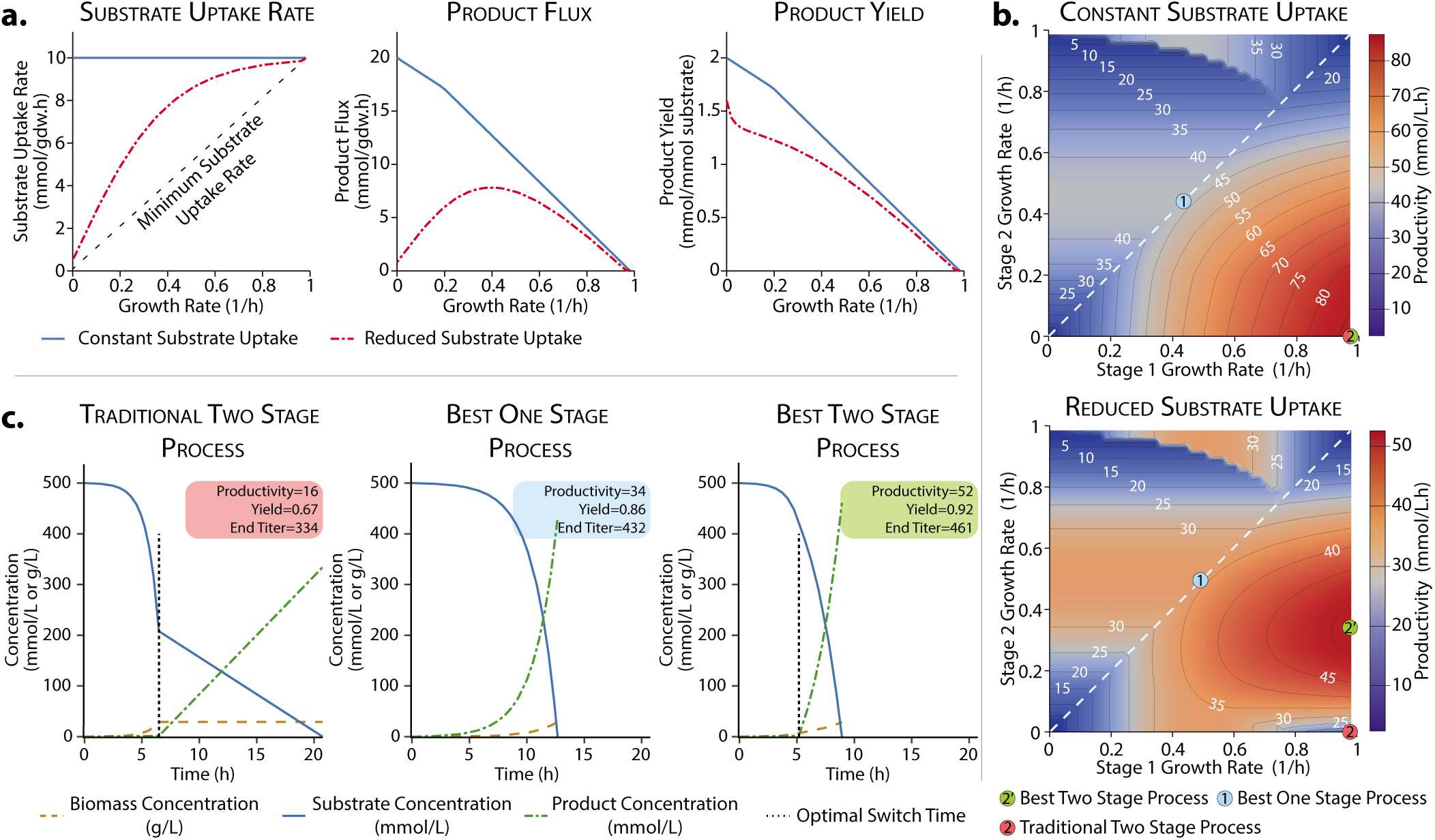
mcPECASO implemented for D-lactic acid production in *E. coli*. *Objective I* - maximizing productivity **a.** substrate uptake rates, product flux and product yields obtained using the iJO1366 reconstruction of *E. coli* assuming constant and reduced substrate uptake rates. In the panel for substrate uptake rate, the minimum required substrate uptake rate at each value of growth rate (shown using a black dashed line) was obtained by performing flux balance analysis on the metabolic model by setting substrate uptake as the objective. **b.** Productivity distribution for TS and OS processes in *E. coli* assuming constant and reduced substrate uptake rates. Isoclines on the distributions show phenotypes with the same productivity levels. **c.** Fermentation profile for various production strategies assuming reduced substrate uptake rates.

However, an impediment in growth rate either due to reaching stationary phase or rewiring of metabolism has been shown to alter substrate uptake rates^25–31^. If these effects are considered, the product flux in a non-growing strain is heavily impacted (shown as red dashed lines in Figure 3a). This makes a nongrowing phenotype during the second stage of a TS process ineffective. We can observe this in Figures 3b,c where the traditional TS process has very low productivity - among the lowest of any possible process. As observed in a previous study^32^, many OS processes (represented on the dashed lines in Figure 3b where the growth and production phenotypes are the same) have higher productivity than the traditional TS process. However, a fair evaluation of two-stage strategies should include the entire available phenotypic space. Even under reduced substrate uptake conditions, there are several TS processes that have higher productivity than the best performing OS process. These processes can be achieved by allowing the strain to grow at a reduced rate during the production stage, rather than completely eliminating growth. The switch time optimization formulation results in earlier switching between the phenotypes when the strain is allowed to grow during the production stage (Figure 3c. The TS process with the highest productivity requires the strain to be able to dynamically switch from wild-type growth to a phenotype with intermediate growth and production (growth coupled production). A sensitivity analysis performed on the bioprocess objective’s response to varying the initial substrate and biomass concentration showed that TS processes outperformed OS processes over a broad range of fermentation start conditions (Supplementary Figures S4 and S5).

However, a process with the highest productivity may not always be the economically optimal choice due to variations in substrate and product prices. *Objective I* aims to maximize productivity with no constraints on the yield resulting from the process. Hence, to analyze TS and OS processes in a more realistic manner, we examined each process type with two additional objectives - *Objective II* which maximizes productivity with an added requirement that the yield must be at least 75% of the maximum value and *Objective III* which maximizes yield with a minimum productivity of 2 *g/L.h*. These values have been previously cited to be the minimum acceptable values for a bioprocess that targets commodity chemicals to be economically viable^15^. We find that the best TS process results in higher productivity than all OS alternatives in the case of *Objective II* as well (Supplementary Figures S2a and S3a) We also examined the effect of varying the minimum required yield on the productivity achieved by the various process types and found that TS processes result in significantly higher productivity over the entire range of possible yield constraints (Supplementary Figure S6a). For *Objective III*, best performing OS process seems to result in the highest yield (Supplementary Figures S2b and S3b). However, at higher requirements of productivity, the yield from the OS process becomes lower than the TS process and above a certain threshold, the OS process becomes infeasible since it is not possible to satisfy the productivity constraint. This shows that TS processes outperform OS processes for D-lactate production in *E. coli* over a broad range of requirements and fermentation start conditions if a switch from wild-type growth to an intermediate growth coupled production phenotype is made.

### 3.2 TS processes are optimal for all natural metabolites in *E. coli*

Having established that TS processes outperform OS alternatives for lactic acid production using economically relevant objectives, we wished to examine if they could dominate OS processes for other natively produced metabolites in *E. coli* for all three objectives. We anticipated that the different production flux profiles for each product would result in variations in the process metrics. Moreover, the constraints in two of the bioprocess objectives could result in different processes being more suitable for each product. Hence, we used mcPECASO with the fermentation start parameters previously described, to predict process optimality for 70 native exchange metabolites (metabolites that appear in exchange reactions) in the iJO1366 reconstruction of *E. coli* ‘s metabolism. For *Objective 1* and *Objective II* where the goal is to maximize productivity, the best TS processes have the highest productivity for all products analyzed, with the OS process and traditional TS process trailing behind (Figure 4a and Supplementary Figure S7a). In general, products with more carbon atoms have lower molar productivity. However, two products, namely 5-methylthioribose and spermidine have unusually low productivities, which will be examined in later sections. The traditional TS processes have very stunted productivities for all the exchange metabolites when substrate uptake rates is reduced, consistent with the previous study for D-lactic acid^32^.

**Figure 4:**
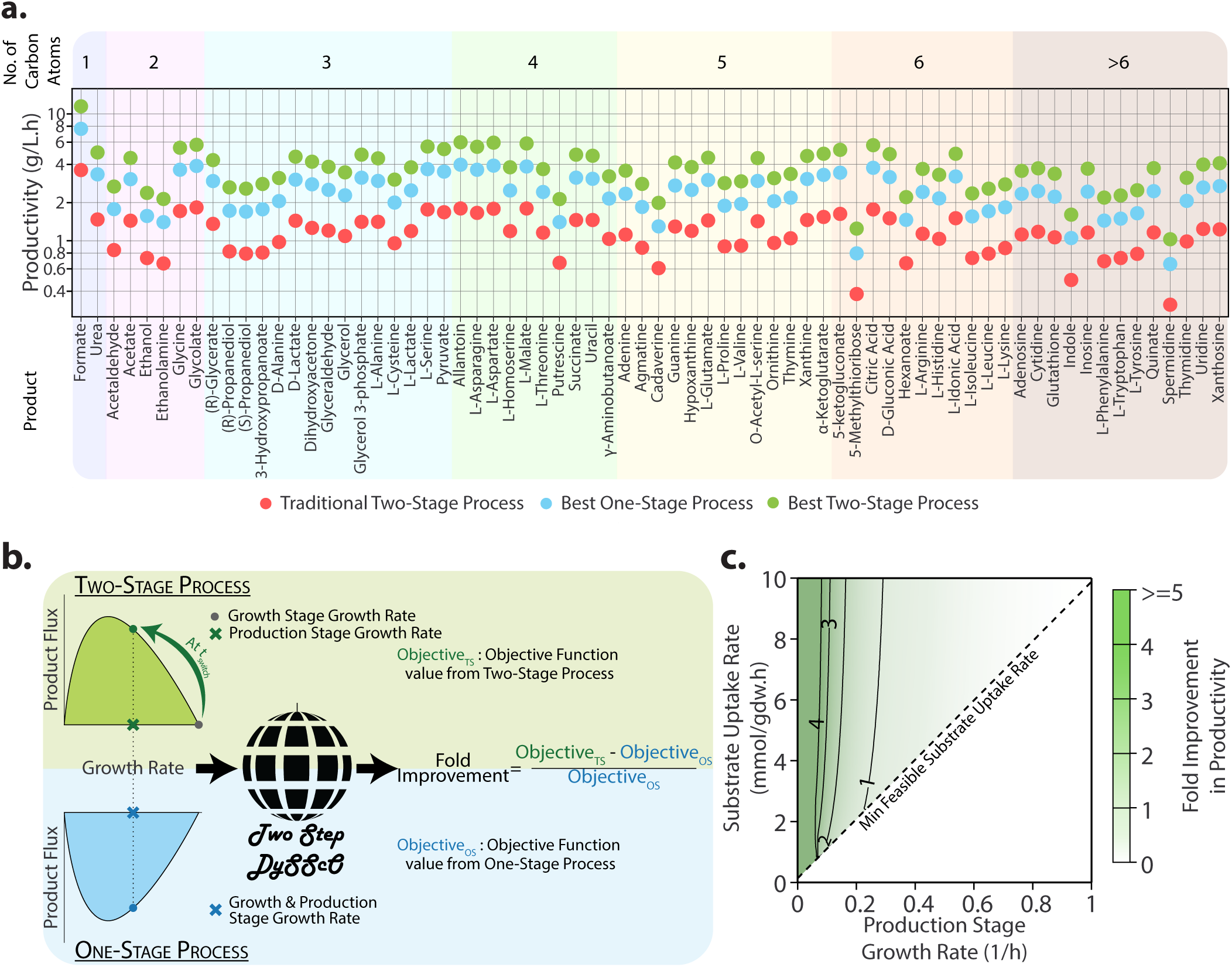
Comparison of TS processes for chemical production using *E. coli*. **a.** Productivity for exchange metabolite production in *E. coli*. mcPECASO was used to simulate the production of all exchange metabolites for *Objective I* - maximizing productivity in the iJO1366 reconstruction of *E. coli*, ordered by the number of carbon atoms in the product. The optimized two stage process always results in the highest productivity. **b.** Workflow to compare TS and OS processes with different substrate uptake characteristics. Production stage phenotypes for TS and OS processes are compared by computing the fold improvement in the objective function resulting from either process type. **c.** Fold improvement in productivity when using a TS process with *Objective I* - maximizing productivity for native exchange metabolites in *E. coli*. TS processes result in higher productivity at all points in the figure, indicating the superiority of such processes regardless of the substrate uptake rate prevailing during the production stage.

Surprisingly, when feasible, TS processes result in higher yields than OS processes when the objective is to maximize process yield with a minimum required productivity (*Objective III*) as seen in Supplementary Figure S7b. This is due to OS processes with higher yields being unable to satisfy the productivity constraint specified by *Objective III*. For the lone case of D-lactic acid, it appears that an OS process with the same yield as a TS process is able to satisfy the productivity requirement. All three process types are infeasible for three products - Spermidine, Indole, and 5-Methylthioribose. For all other products where productivity and yield requirements are feasible, TS processes dominate. OS processes even become infeasible, being unable to satisfy the productivity constraint for many products. Hence, TS processes are of value even when yield maximization is the objective in an industrial setting.

Upon examining the best TS processes predicted by mcPECASO, we found that all of them had the same phenotype during the growth stage - wild-type growth. These processes varied in phenotype only during the production stage, where an intermediate growth phenotype resulted in the highest productivity. Similarly, OS processes with the highest productivity are those with an intermediate growth rate and have one growth-coupled production stage. Hence, it is possible to compare the two process types at every operating point in the production envelope during the production stage for all possible substrate uptake rates, allowing us to come to more generalizable conclusions. As stated before, the logistic curve used to model substrate uptake variation is just an approximation that seeks to closely model a worst-case scenario for substrate uptake rates. This analysis where we allow substrate uptake to take any value above the minimum feasible rate would help in establishing the superiority of one process type over another.

To compare the process types for the three different objectives, we calculated the fold improvement in the objective value offered by the TS process when compared to the OS process at each feasible operating point (Figure 4b). For *Objective I* and *Objective II*, we found that the fold improvement in productivity remained the same for all 70 products. TS processes have higher productivity at all feasible substrate uptake rates for *Objective I* (Figure *4c). In the case of Objective II*, an increase in the yield constraint reduces the range of substrate uptake rates over which either process is feasible (shown by the red region in Supplementary Figure S8). However, TS processes still result in higher productivity over the entire feasible region of substrate uptake rates.

However, for *Objective III*, each product resulted in a different distribution of the fold improvement in process yield. This is because the constraint on productivity is given in mass units and not molar units. Regardless, we analyzed each product individually and have presented them in Supplementary Figure S9. Apart from a few exceptions, OS processes are not feasible for most products over a large range of substrate uptake rates.They are feasible only when the production stage growth rates and the corresponding substrate uptake rates are high. At the operating points where feasible, they result in higher yields. However, TS processes are optimal over a larger range of substrate uptake rates for most products and are able to achieve the same yields as OS processes at lower growth rates. Moreover, for every product, the highest yield obtainable using TS processes is more than the highest yield possible using OS processes, as indicated by contour lines in each plot of Supplementary Figure S9. Hence, TS processes are optimal for yield and productivity maximization, irrespective of the substrate uptake characteristics.

### 3.3 Similarities in production stage fluxes suggest the possibility of a universal production phenotype

While it is clear that the enhanced TS processes obtained here are optimal for maximizing yield and productivity, it would be useful to determine how such processes can be physically realized. Hence, we examined the flux perturbations required to achieve the various production strategies for each of the exchange metabolites analyzed in the previous section using flux variability analysis, as described previously. The best TS processes predicted by mcPECASO use wild-type growth as the phenotype in the first stage. Therefore, using the wild-type flux distribution as a reference, we examined the number of reactions that would need to be perturbed for each product in the three objectives (Supplementary Figures S10, S11, and S12), and classified each perturbed reaction/flux them based on whether they are:

- Switched On - growth stage flux is zero and production stage flux is non-zero.
- Switched Off - growth stage flux is non-zero, production stage flux is zero.
- Significantly Upregulated - production stage flux is at least 10% more than the growth stage flux
- Significantly Downregulated - production stage flux is decreased more than biomass flux.

Fewer than 10 reactions need to be switched on or completely switched off for most products in all three objectives to achieve the production stage. Products with fewer carbon atoms require more reactions to be turned on/off and consequently, a bigger flux change compared to larger products. This suggests that most of the reactions required to produce larger products optimally are expressed during wild-type growth. Very few reactions need to be completely switched off for most products, implying that reactions involved in biomass synthesis are required in some capacity during the production stage as well. Interestingly, the compounds that were determined to have an unusually low yield in the previous section - 5-methylthioribose and spermidine have among the largest number of significantly upregulated reactions of all products. The yield of these products is likely low due to the upregulation of pathways that result in yield losses during production and the absence of alternative pathways that conserve yield.

To further understand the production-stage phenotypes, we examined the magnitude of flux changes from the wild-type flux distribution for every product and arranged the reactions based on the subsystem in which they occur for each objective (Figure 5 and Supplementary Figure S13). Most perturbations for the best TS process are downregulations, with a majority of these reactions having the same level of reduction in flux as the reduction growth rate (about 60%). Hence most of the flux changes observed are effected by a decrease in growth/biomass synthesis rate. This is also evident from the fact that all reactions involved in membrane lipid metabolism and cell envelope biosynthesis are consistently downregulated for most products. Similarly, glycolysis and citric acid cycle (TCA cycle) pathway reactions are downregulated for most products. However, many reactions involved in glycolysis have a much smaller change, indicating that carbon flow remains consistent through these reactions and flux is partitioned towards production pathways further downstream. Interestingly, reactions in the pentose phosphate pathway are equally divided between being upregulated or downregulated together for different products. Hence, this subsystem appears to act as a key node that controls precursor and cofactor availability to manufacture metabolites optimally within the cell. These results suggest that it is possible to engineer an *E. coli* strain with a universal production phenotype that maximizes productivity/yield. In such a strain, flux perturbations that appear for all the products of interest can be dynamically implemented by throttling flux through key reactions contributing to biomass synthesis. Then, depending on the class of product of interest - amino acid, nucleotide, central carbon metabolite, etc, those reactions/pathways that require upregulation can be set up in a modular manner and dynamically expressed if required.

**Figure 5:**
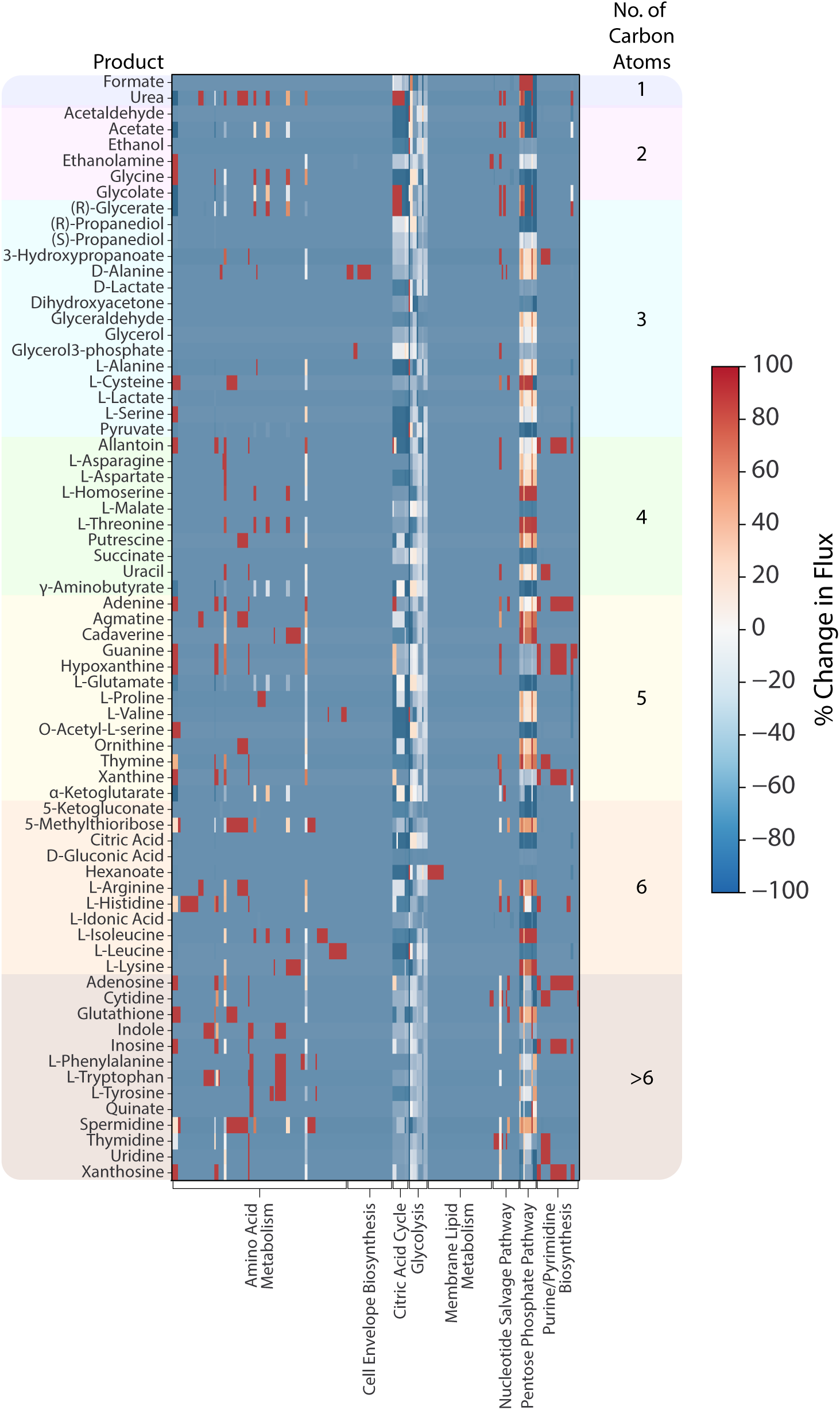
Flux perturbations required for TS processes. Percent change in flux through all reactions compared to wild-type flux distribution required to achieve the TS processes for native exchange metabolites in *E. coli* for *Objective I* - maximizing productivity. Due to the large number of reactions involved, only the reaction subsystems are shown. A large number of reactions show a 60% reduction in flux for all products, which is caused by an identical decrease in growth rate. Flux changes that are not effected by growth rate reduction are mostly localized to specific reaction subsystems.

### 3.4 Perturbations increasing phosphoenolpyruvate and NADPH availability are enriched

In order to identify key control reactions, we analyzed which reactions are enriched in production strategies for the exchange metabolites previously analyzed. We did this by looking at the number of products for which each reaction appeared as a perturbation and classified them based on the type of perturbation - on, off, upregulated or downregulated for each objective (Figure 6 and Supplementary Figure S14). Only non-transport reactions involved directly in metabolism were retained for the final analysis. The full names of reactions and their corresponding reaction formulae are available in Supplementary Table S1.

**Figure 6:**
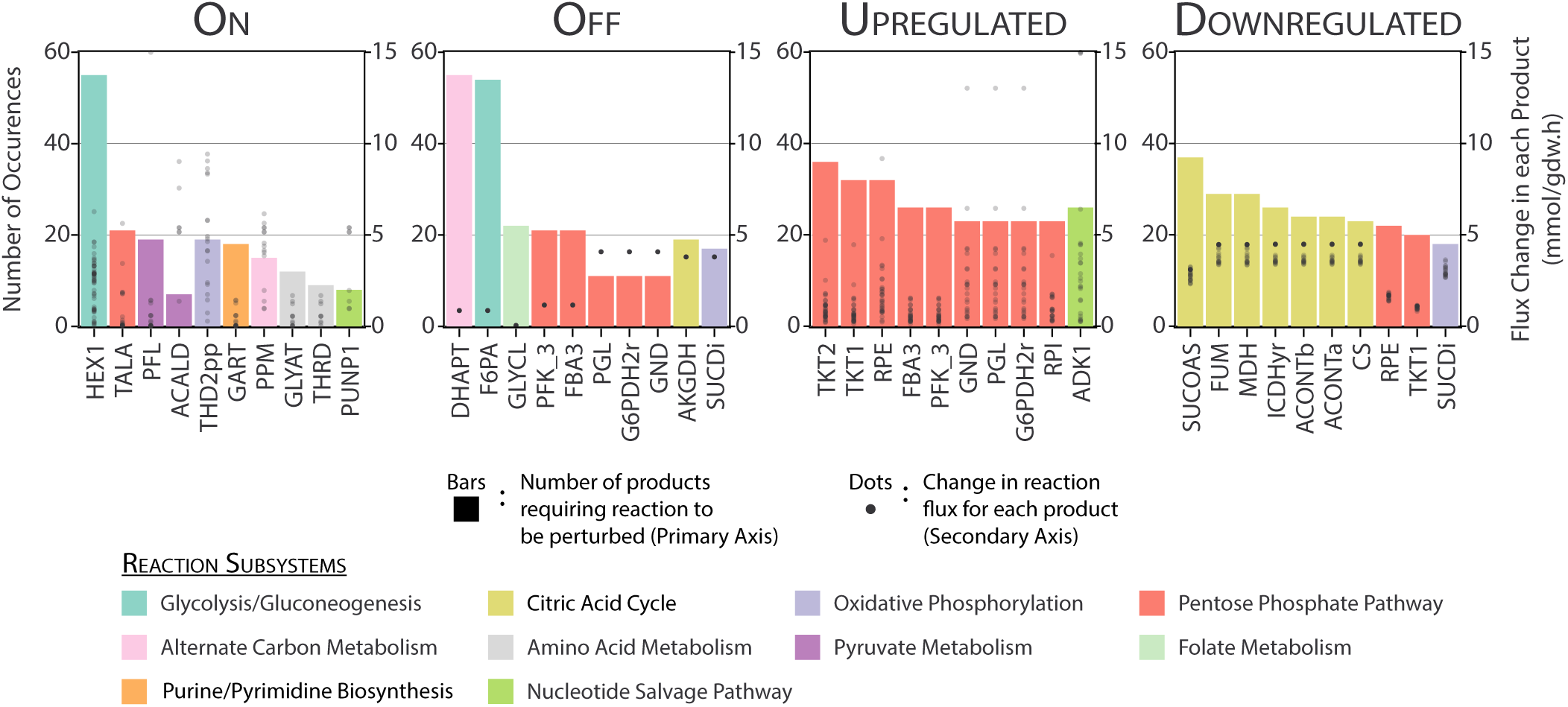
Most frequently occurring reactions in perturbations for TS processes in *E. coli*. Top ten frequently occuring reactions in the two-stage production strategies for native exchange metabolites in *E. coli* for *Objective I* - maximizing productivity, were obtained and classified based on the type of perturbation. The corresponding change in fluxes required in the reactions for each product are represented as dots with values in the secondary axis.

Among the reactions that are switched on for TS strategies (Figure 6), HEX1 (ATP dependent hexokinase) occurs in more than 75% of the products. This reaction serves as an alternative to the phosphotransferase system that is used by wild-type *E. coli* to transport and phosphorylate glucose. It differs from the conventional phosphotransferase system, in that it uses ATP for phosphorylation as opposed to phosphoenolpyruvate (PEP). PEP is a key precursor to many products and therefore, alternative means of glucose usage that use less PEP are enriched in the ‘on’ type perturbations. Interestingly, the HEX1 glucose transport system has previously been studied for its importance in creating platform strains for microbial chemical production^50^. Furthermore, extra usage of ATP has been explored as a means to increase substrate uptake rates, potentially increasing product flux under low growth conditions^51^, suggesting the importance of studying alternatives to the native phosphotransferase system. The reaction THD2pp (NADP transhydrogenase) is also required to be switched on for many products. This likely serves to increase NADPH availability to cater to production pathways.

Among other types of perturbations, DHAPT (Dihydroxyacetone phosphotransferase) and F6PA (Fructose 6-phosphate aldolase) are reactions that need to be turned off for most products. These also likely serve to collectively increase PEP availability for production reactions since DHAPT utilizes PEP. Many upregulated reactions for *Objective I* are from the pentose phosphate pathway subsystem, serving to increase the availability of NADPH and pentose sugars for products. Surprisingly, apart from the transketolase (TKT2), none of the other reactions in the pentose phosphate pathway seem to be significantly upregulated for the other two objectives. However, the overall number of upregulations is also lower in these cases. TCA Cycle (Citric acid cycle) reactions are most frequently downregulated for all three objectives.

Notably, with the exception of the pentose phosphate pathway reactions in *Objective II* and *Objective III*, the same reactions occur as frequent perturbations for all three objectives. Also, in the case of reactions that are switched off/downregulated, some are sequential reactions({G6PDH2r, PGL, GND}, {FUM, MDH}, etc)l and need not all be downregulated/switched off to control flux through that pathway. Moreover, most of these reactions require very similar changes in their fluxes (shown as dots in Figure 6 and Supplementary Figure S14) for the products in which they occur. These features lend credence to the possibility of the universal production strain discussed in the previous section strain and make its construction more feasible.

## 4 Conclusions

We have seen that the choice of process type influences the process metrics and therefore the profitability of a microbial chemical production process to a great extent. Furthermore, strain design choices are also influenced by process choice. One-stage processes require static genetic intervention strategies that couple growth and production whereas, two-stage processes require dynamic intervention strategies where gene expression is temporally controlled. Recent advances in CRISPR^52^, transcriptional switches^53^, riboswitches^54^, and other gene regulatory elements present an exciting outlook for the experimental implementation of such intervention strategies. There has also been interest in computational algorithms that predict dynamic control strategies which begin with high growth and switch over to growth-coupled production as required by the best TS production strategies predicted in this study^55^.

Strain engineering efforts necessary to achieve target production phenotypes predicted by mcPECASO may seem daunting due to the number of perturbations required. However, it is important to note that this analysis does not take into account the fact that many pathways are linear and sequential. Therefore, it would not be necessary to actively perturb all fluxes predicted in this study. A reduction of the metabolic network would help to identify key control reactions that actually need to be perturbed. Furthermore, algorithmic approaches can be used to predict the minimal set of genetic perturbations required to achieve target phenotypes given the constraints predicted by mcPECASO^56^. Nevertheless, even with a large list of candidate genes, it is possible to use a combination of high-throughput experiments, cell sorting and a rational subset of candidates to identify which perturbations that actually lead to an improvement in production characteristics, as seen in a recent study^57^.

It is interesting to note that the production of most products involves enhancing PEP conserving and NADPH overproducing strategies. The emergence of PEP as a key precursor indicates its importance as a bow-tie metabolite, funneling flux into different pathways^2,58^. Furthermore, reactions in the pentose phosphate pathway seem to work in unison to increase precursor availability for number of products by being upregulated or downregulated, alluding to their importance in making metabolic networks malleable and robust to perturbations. These common features in strains with enhanced production of a wide range of metabolites give rise to the idea of a universal production strain that could be used to maximize productivity in a TS process by redirecting flux from growth to production related processes for various classes of products. Such a platform strain that maximizes productivity could be realized by placing a minimal number of control reactions under dynamically repressible/inducible promoters to throttle biomass production flux. Tools that allow the dynamic perturbation of a large number of genes simultaneously could hold the key to realizing such strains^52^. This is similar to the concept of a modular cellular chassis for the production of many different compounds, that has gained interest recently^59,60^.

Previous studies have only analyzed TS processes for their importance in improving process productivity. Here, we have shown that TS processes are able improve both process productivity and overall yield for native exchange metabolites in *E. coli* even if the substrate uptake limitation caused by reduced growth is considered. Since the production characteristics can be expected to be similar for other organisms and non-native products too, this conclusion can be extended to products in other hosts as well. We found that this conclusion holds true over a wide range of industrially relevant fermentation start parameters. In future, better substrate uptake rate measurements for strains with production phenotypes will help in making more accurate predictions of process performance. While it is true that the process metrics depend on the substrate uptake rate of the mutant strain, we have shown that a TS process can outperform an OS process, irrespective of substrate uptake characteristics for all economically relevant bioprocess objectives. Further improvements in substrate uptake rates through various strategies^61^ will improve process metrics even further. The software framework presented here - mcPECASO, has the ability to determine the effectiveness of each process type and predict optimal hypothetical phenotypes for experimental evaluation. It also provides information about the fermentation conditions under which each process type would perform better. We anticipate that mcPECASO and the findings obtained in this study will be very valuable to make process and strain design decisions for industrial scale production of chemicals using microorganisms. Furthermore, the concept of a universal production strain that has the same growth phenotype and several common flux perturbations identified here to switch to a production phenotype for a diverse range of chemicals in a flexible manner may provide a paradigm shift in the way chemical production processes are designed in the future.

## Supporting information

Experimental substrate uptake data

## Acknowledgements

The authors thank Ruhi Choudhary (University of Toronto) for suggesting structural edits to the manuscript and Kevin Correia (University of Toronto) for insightful discussions regarding the optimization framework of mcPECASO.

## Author’s contributions

KR helped formulate the study, contributed to the codebase, implemented the framework, and wrote the article. NV helped formulate the study, contributed to the codebase, and edited the manuscript. RM helped formulate the study, supervised the work, and edited the manuscript.

