## Supplementary methods, tables and figures for "Novel two-stage processes for optimal chemical production in microbes"

### Supplementary Information

#### 1 Methods

##### 1.1 Approximating substrate uptake rate variation in *E. coli*

Figure S1a shows glucose uptake rate data obtained from experiments on various strains of *Escherichia coli*. Since strains from different backgrounds can be expected to exhibit different phenotypes upon genetic perturbations, separate models need to be developed for each of these. Growth data for different strains were obtained from literature and have been compiled in the supplementary data file **'experimental_substrate_uptake_data.xlsx'** We only consider one strain backgrounds - MG1655 for modeling the effects of growth rate on substrate uptake due to the availability of a large number of experimental datapoints and a well curated metabolic model for use in this study. It is also important to note that the large errors in the measured growth and substrate uptake rates preclude the ability to develop an accurate model for substrate uptake variation. We fit the experimental data to a linear equation and simplified logistic curve. Though the straight line fit has a lower $\chi^2$ value than the logistic curve fit, we expect that the logistic equation represents a more realistic model of substrate uptake rate variation with graded changes in phenotypes (Figure S1b). This is supported by studies which show that phosphotransferase activity and correspondingly, glucose uptake rates reduce in *E. coli* with increasing concentrations of allosteric regulators including $\alpha$-ketoglutarate and oxoaloacetate, following inhibition kinetics. An additional assumption we use is that as growth rates are reduced in *E. coli*, the levels of intracellular metabolites, including those that inhibit glucose uptake increases. Hence, a saturation type model can be used to approximate glucose uptake variation with growth rate. We chose to use the logistic curve (Eq. S1) since it is the most widely used saturation type model and offers the flexibility of being able to choose start and end points. We chose parameter values such that the model is able to accurately predict experimentally observed substrate uptake values at stationary phase and wild-type growth, while giving the minimum possible substrate uptake rate at intermediate growth rates.

$$
\begin{aligned}
q_s &= q_{s,min} + (q_{s,max} - q_{s,min}) * \left(-1 + \frac{2}{(1 + e^{-K_{uptake} * \mu})}\right) \\
&\text{where} \\
q_{s,min} &= 0.5\ mmol/gdw.h \\
q_{s,max} &= 10\ mmol/gdw.h \\
K_{uptake} &= 5
\end{aligned}
\tag{Eq. S1}
$$

### 2 Supplementary tables

**Table S1:** Names and formulae of reactions mentioned mentioned in Figure 6 (main text) and Figure S14

| Reaction ID | Reaction Name | Reaction Formula |
|---|---|---|
| ACALD | Acetaldehyde dehydrogenase | acald_c + coa_c + nad_c <=>accoa_c + h_c + nadh_c |
| ACONTa | Aconitase A | cit_c <=>acon_C_c + h2o_c |
| ACONTb | Aconitase B | acon_C_c + h2o_c <=>icit_c |
| ADK1 | Adenylate kinase | amp_c + atp_c <=>2 adp_c |
| AKGDH | 2-Oxogluterate dehydrogenase | akg_c + coa_c + nad_c ->co2_c + nadh_c + succoa_c |
| ASPTA | Aspartate transaminase | akg_c + asp__L_c <=>glu__L_c + oaa_c |
| CS | Citrate synthase | accoa_c + h2o_c + oaa_c ->cit_c + coa_c + h_c |
| DHAPT | Dihydroxyacetone phosphotransferase | dha_c + pep_c ->dhap_c + pyr_c |
| F6PA | Fructose 6-phosphate aldolase | f6p_c <=>dha_c + g3p_c |
| FBA3 | S17BP-G3P-lyase | s17bp_c <=>dhap_c + e4p_c |
| FUM | Fumarase | fum_c + h2o_c <=>mal__L_c |
| G6PDH2r | Glucose 6-phosphate dehydrogenase | g6p_c + nadp_c <=>6pgl_c + h_c + nadph_c |
| GART | GAR transformylase-T | atp_c + for_c + gar_c ->adp_c + fgam_c + h_c + pi_c |
| GHMT2r | Glycine hydroxymethyltransferase | ser__L_c + thf_c <=>gly_c + h2o_c + mlthf_c |
| GLYAT | Glycine C-acetyltransferase | accoa_c + gly_c <=>2aobut_c + coa_c |
| GLYCL | Glycine Cleavage System | gly_c + nad_c + thf_c ->co2_c + mlthf_c + nadh_c + nh4_c |
| GLYCL | Glycine Cleavage System | gly_c + nad_c + thf_c –>co2_c + mlthf_c + nadh_c + nh4_c |
| GND | Phosphogluconate dehydrogenase | 6pgc_c + nadp_c ->co2_c + nadph_c + ru5p__D_c |
| HEX1 | Hexokinase | atp_c + glc__D_c ->adp_c + g6p_c + h_c |
| ICDHyr | Isocitrate dehydrogenase | icit_c + nadp_c <=>akg_c + co2_c + nadph_c |
| MDH | Malate dehydrogenase | mal__L_c + nad_c <=>h_c + nadh_c + oaa_c |
| PFK_3 | Phosphofructokinase (s7p) | atp_c + s7p_c ->adp_c + h_c + s17bp_c |

**Table S1:** Names and formulae of reactions mentioned mentioned in Figure 6 (main text) and Figure S14

| Reaction ID | Reaction Name | Reaction Formula |
|---|---|---|
| PFL | Pyruvate formate lyase | coa_c + pyr_c ->accoa_c + for_c |
| PGCD | Phosphoglycerate dehydrogenase | 3pg_c + nad_c –>3php_c + h_c + nadh_c |
| PGL | 6-phosphogluconolactonase | 6pgl_c + h2o_c ->6pgc_c + h_c |
| POR5 | Pyruvate synthase | coa_c + 2 flxso_c + pyr_c <=>accoa_c + co2_c + 2flxr_c + h_c |
| PPC | Phosphoenolpyruvate carboxylase | co2_c + h2o_c + pep_c ->h_c + oaa_c + pi_c |
| PPM | Phosphopentomutase | r1p_c <=>r5p_c |
| PRPPS | Phosphoribosylpyrophosphate synthetase | atp_c + r5p_c <=>amp_c + h_c + prpp_c |
| PSERT | Phosphoserine transaminase | 3php_c + glu__L_c –>akg_c + pser__L_c |
| PSP_L | Phosphoserine phosphatase | h2o_c + pser__L_c –>pi_c + ser__L_c |
| PUNP1 | Purine-nucleoside phosphorylase | adn_c + pi_c <=>ade_c + r1p_c |
| RPE | Ribulose 5-phosphate 3-epimerase | ru5p__D_c <=>xu5p__D_c |
| RPI | Ribose-5-phosphate isomerase | r5p_c <=>ru5p__D_c |
| SUCDi | Succinate dehydrogenase | q8_c + succ_c ->fum_c + q8h2_c |
| SUCOAS | Succinyl-CoA synthetase | atp_c + coa_c + succ_c <=>adp_c + pi_c + succoa_c |
| TALA | Transaldolase | g3p_c + s7p_c <=>e4p_c + f6p_c |
| THD2pp | NADP transhydrogenase | 2 h_p + nadh_c + nadp_c ->2 h_c + nad_c + nadph_c |
| THRD | L-threonine dehydrogenase | nad_c + thr__L_c ->2aobut_c + h_c + nadh_c |
| TKT1 | Transketolase | r5p_c + xu5p__D_c <=>g3p_c + s7p_c |
| TKT2 | Transketolase | e4p_c + xu5p__D_c <=>f6p_c + g3p_c |

#### 3 Supplementary figures

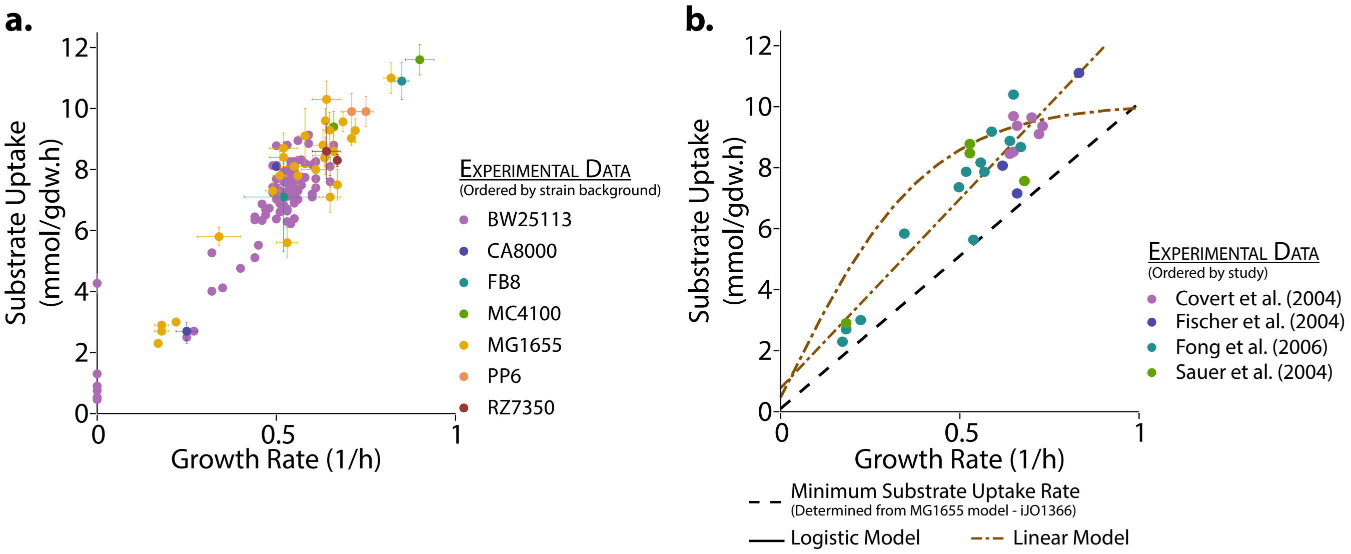


**Figure S1:** **a.** Substrate uptake variation with growth rate for different *E.coli* strains. **b.** Linear and Logistic equation fit for MG1655 experimental substrate uptake data.

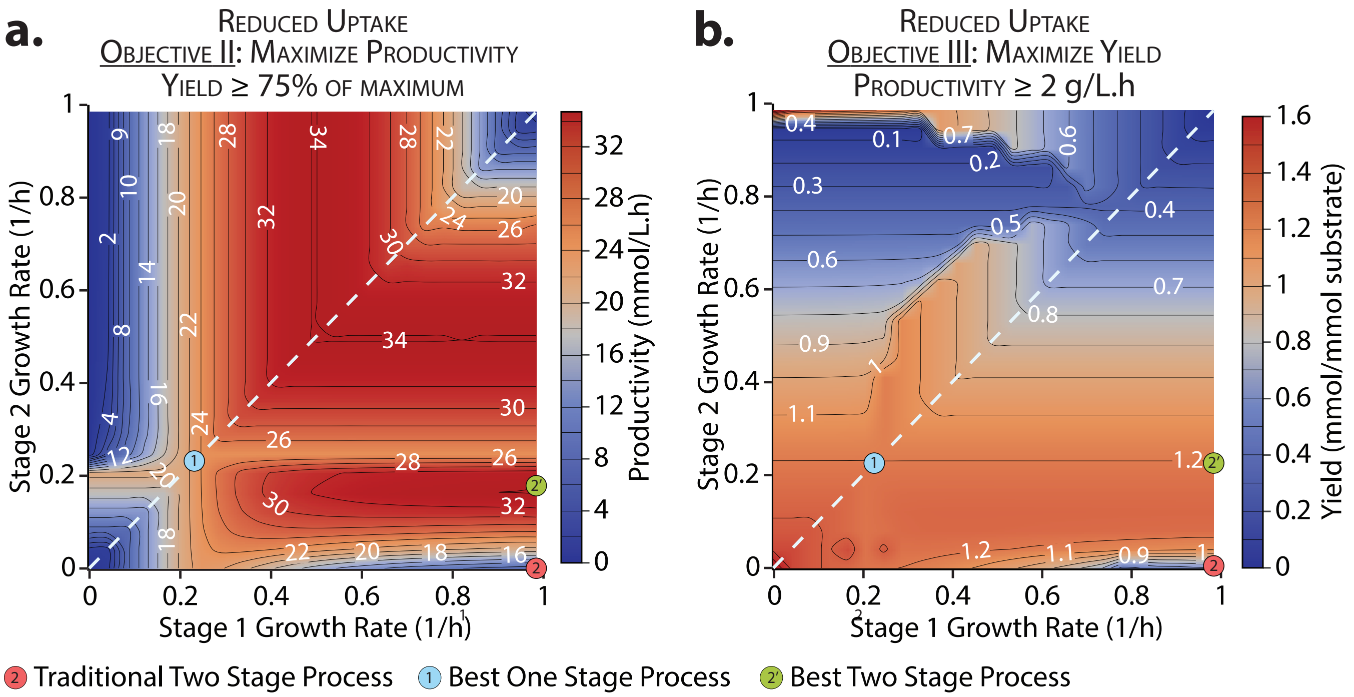


**Figure S2:** Objective value distribution for D-lactic acid production in *E. coli* with $S_0$ (initial substrate concentration) $= 500\,mM$ and $X_0$ (initial biomass concentration) $= 0.05\,g/L$ using different substrate uptake assumptions and fermentation objectives: **a.** Reduced substrate uptake rates and Objective II - maximize productivity with a yield of at least 75% of the maximum value. **b.** Reduced substrate uptake rates and Objective III - maximize yield with a productivity of at least 2 g/L.h.

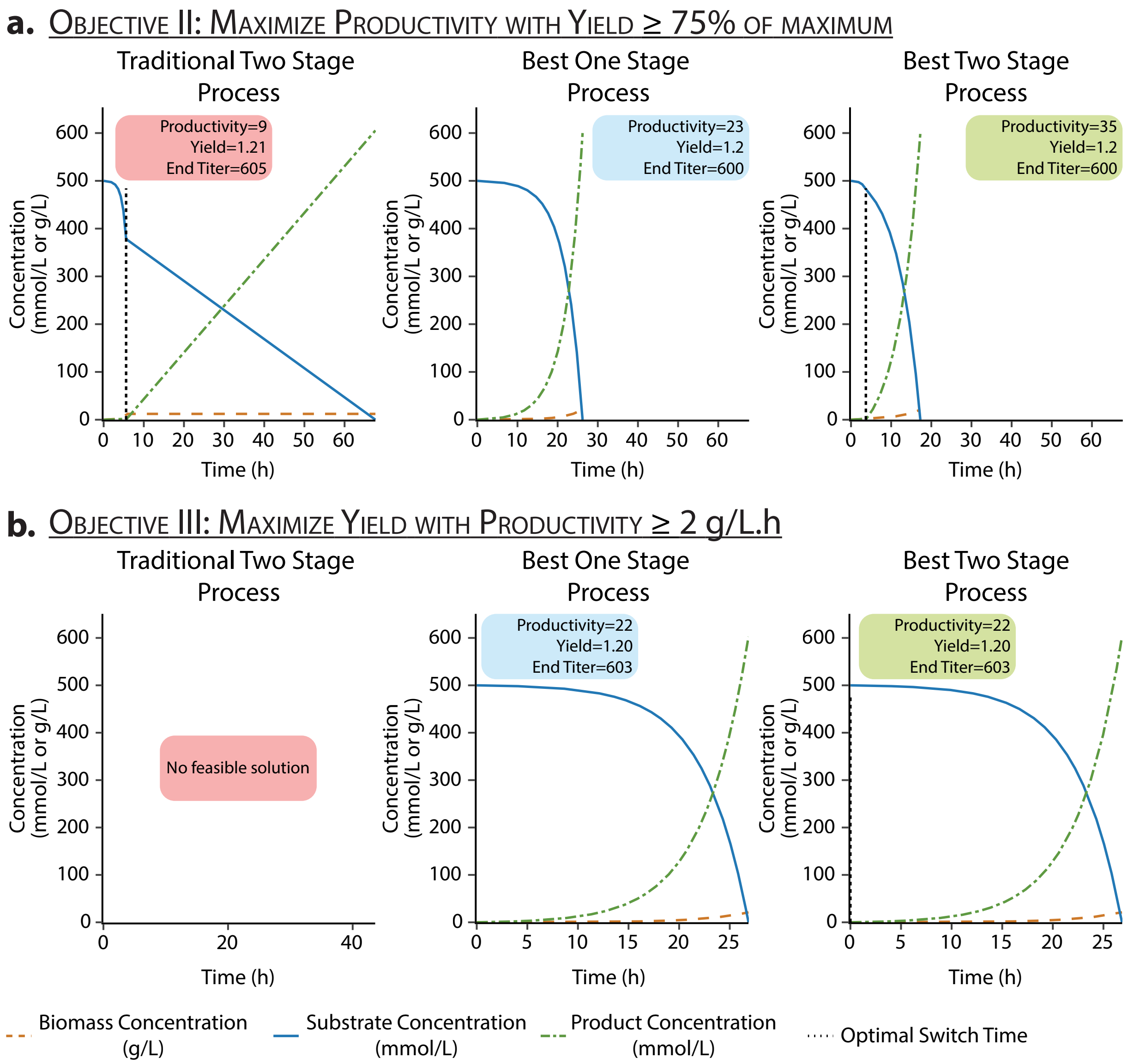


**Figure S3:** Fermentation profile for D-lactic acid production in *E. coli* with $S_0$ (initial substrate concentration) $= 500\,mM$ and $X_0$ (initial biomass concentration) $= 0.05\,g/L$ with different production strategies assuming constant substrate uptake rates and two different fermentation objectives: **a.** Objective II - maximize productivity with a yield of at least 75% of the maximum value. **b.** Objective III - maximize yield with a productivity of at least 2 g/L.h. Note that no feasible traditional two-stage process could be obtained satisfying the given constraints for Objective III.

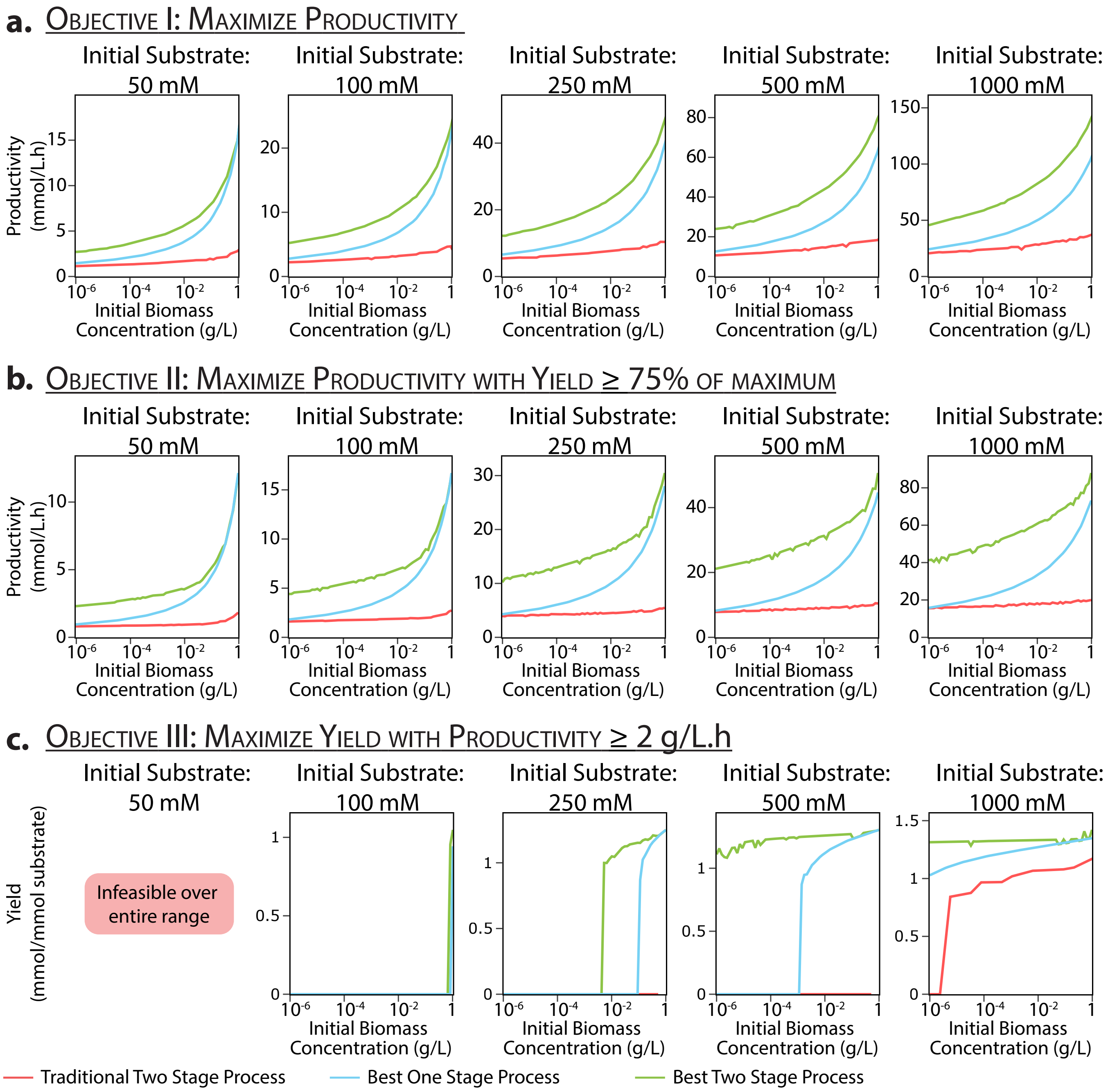


**Figure S4:** Senstivity of objective values to initial biomass concentrations at various initial substrate concentrations for D-lactic acid production in *E. coli* for three different objectives: **a.** Objective I - maximize productivity. **b.** Objective II - maximize productivity with a yield of at least 75% of the maximum value. **c.** Objective III - maximize yield with a productivity of at least 2 g/L.h.

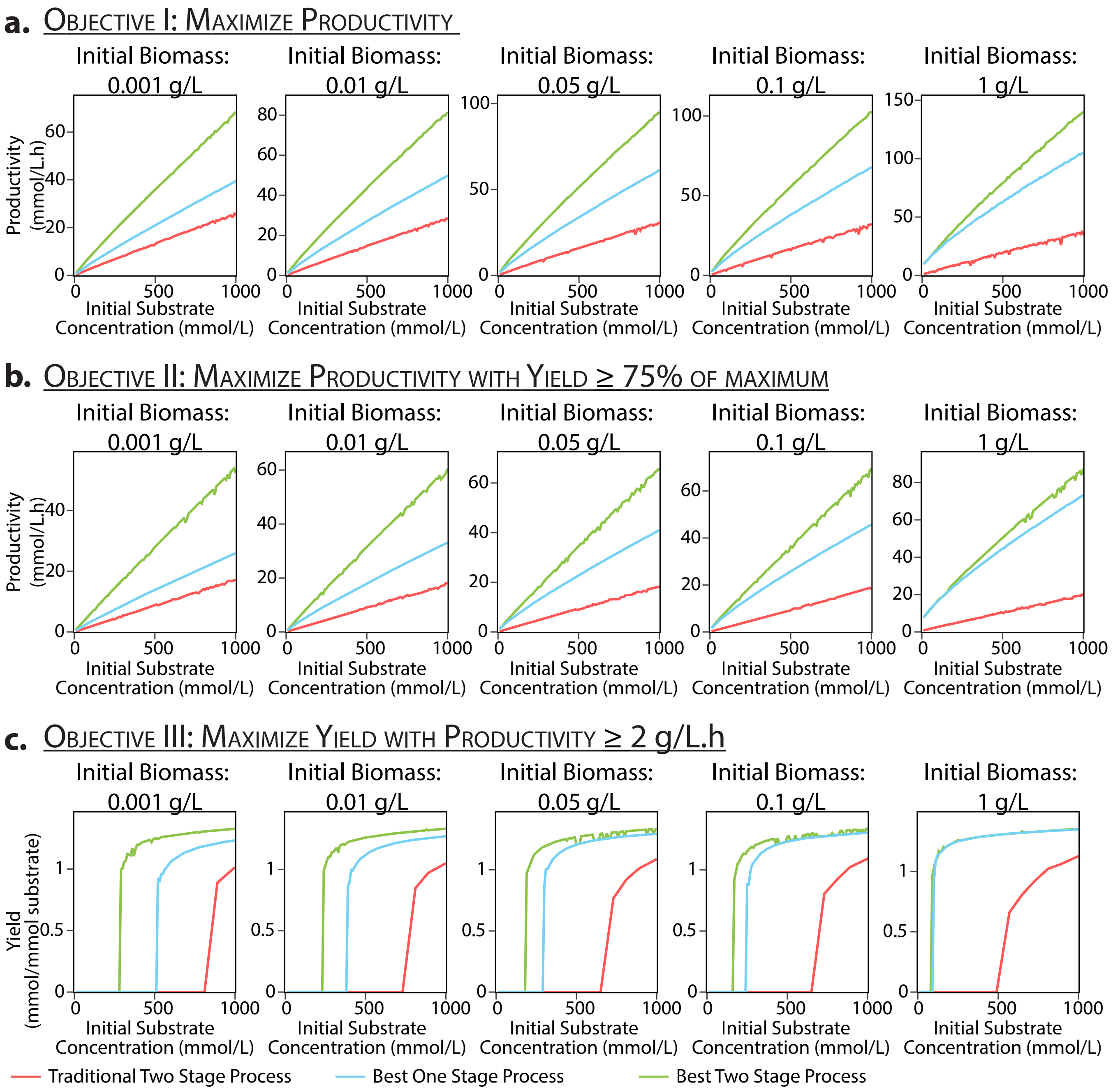


**Figure S5:** Senstivity of objective values to initial substrate concentrations at various initial biomass concentrations for D-lactic acid production in *E. coli* for three different objectives: **a.** Objective I - maximize productivity. **b.** Objective II - maximize productivity with a yield of at least 75% of the maximum value. **c.** Objective III - maximize yield with a productivity of at least 2 g/L.h.

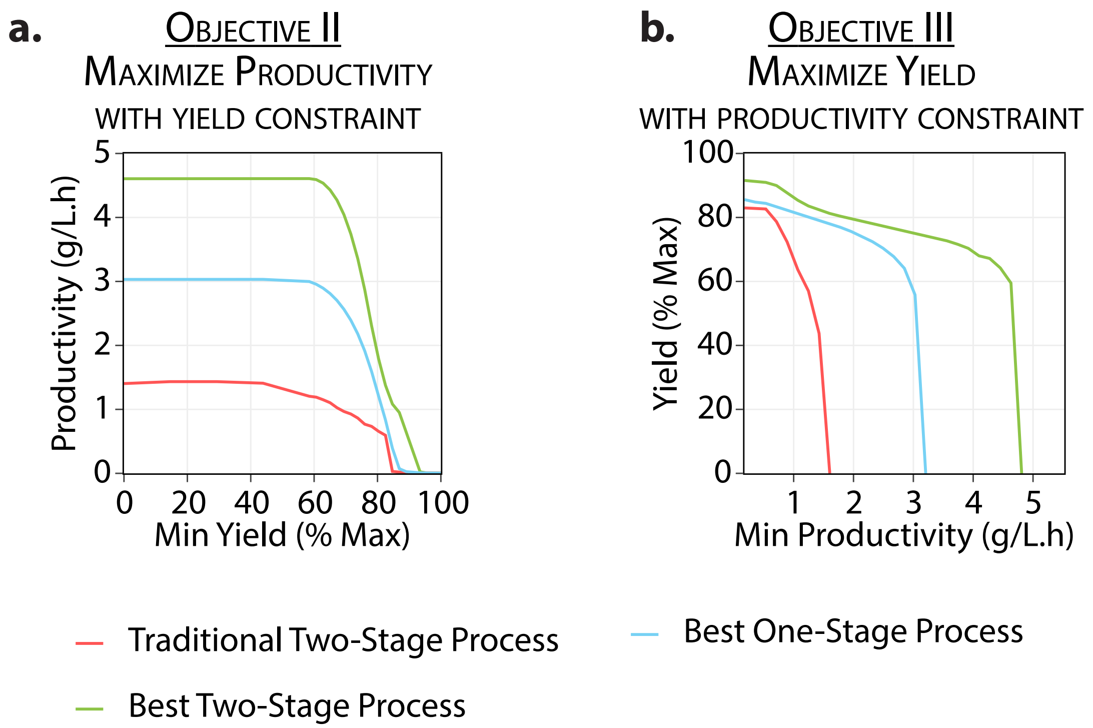


**Figure S6:** Effect of varying fermentation metric constraints on objective values for D-lactic acid production in *E. coli* with **a.**Objective II - Maximize productivity using a varying yield constraint. **b.**Objective III - Maximize yield using a varying productivity constraint.

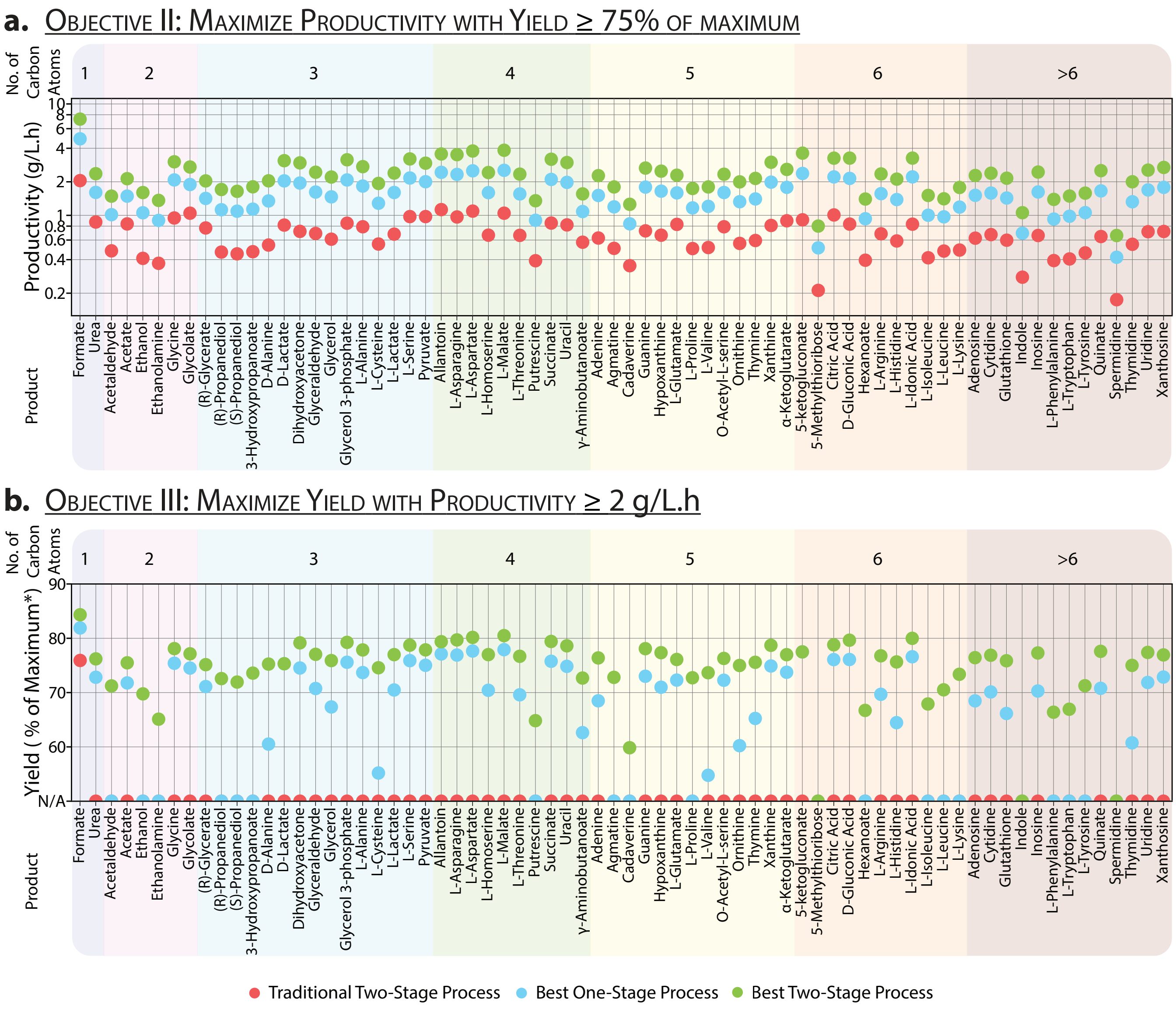


**Figure S7:** Fermentation objective values for native exchange metabolites in *E. coli* using different fermentation strategies with different objectives: **a.** Objective II - maximize productivity with a yield of at least 75% of the maximum value. **b.** Objective III - maximize yield with a productivity of at least 2 g/L.h. Note that for Objective III, an objective value of $'N/A'$ implies no feasible solution satisfying the constraints could be found.

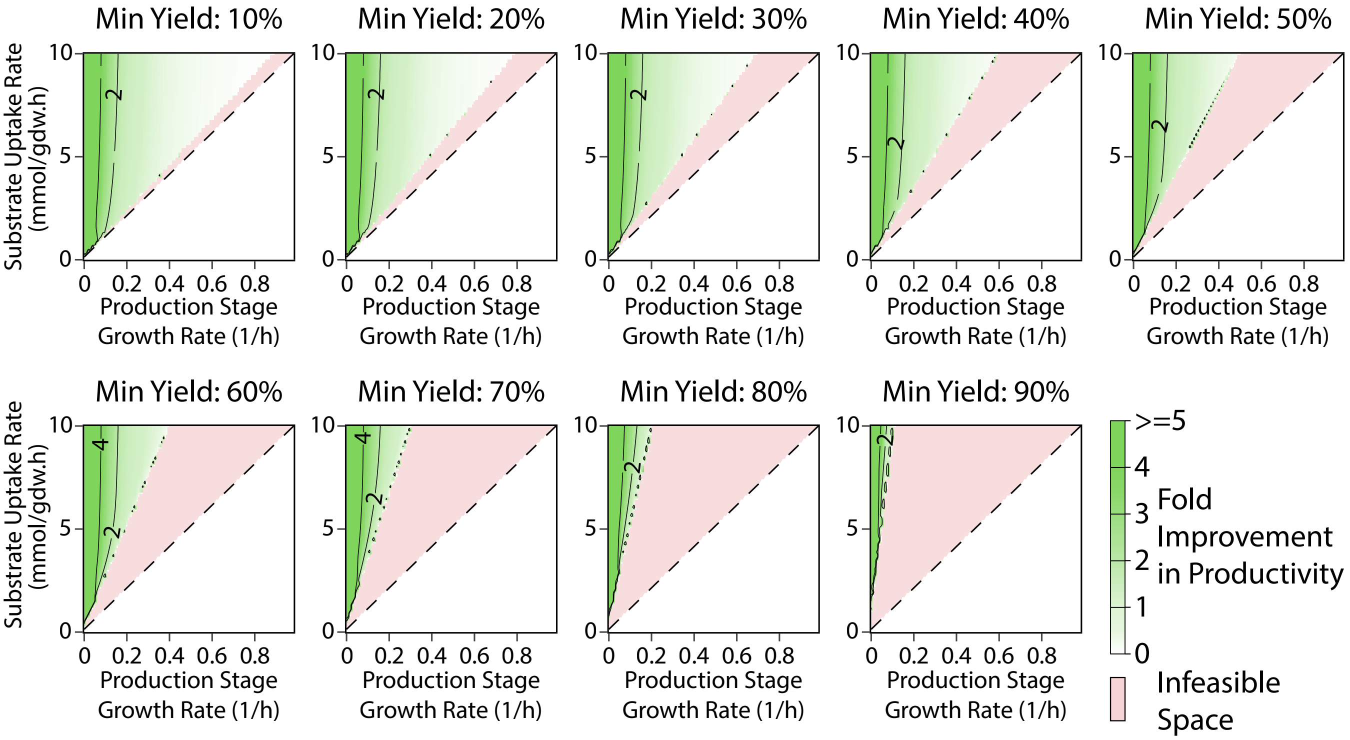


**Figure S8:** Fold improvement in productivity for all native exchange metabolites in *E. coli* from a TS process compared to an OS process at all possible substrate uptake rates during the production stage with fermentation objective II - maximize productivity with varying constraints on yield.

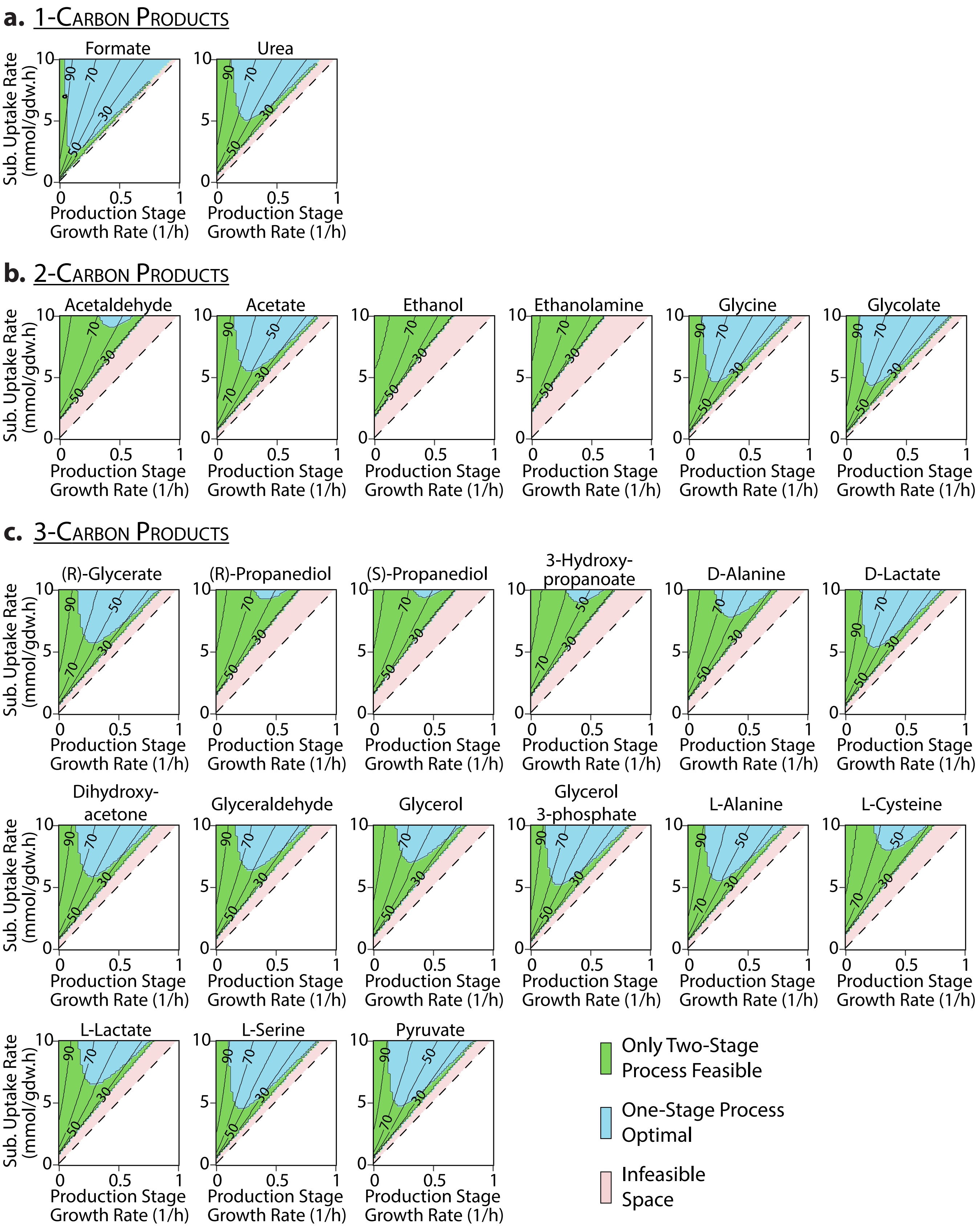


**Figure S9:** Comparison of TS and OS processes in *E. coli* at all possible substrate uptake rates during the production stage with fermentation objective III - maximize yield with a minimum productivity of 2 g/L.h for: **a.** 1 - carbon products. **b.** 2 - carbon products. **c.** 3 - carbon products. The yields at each point within the feasible regions for TS and OS processes are represented by the contour lines as a % of the maximum yield.

#### **d.** 4-Carbon Products

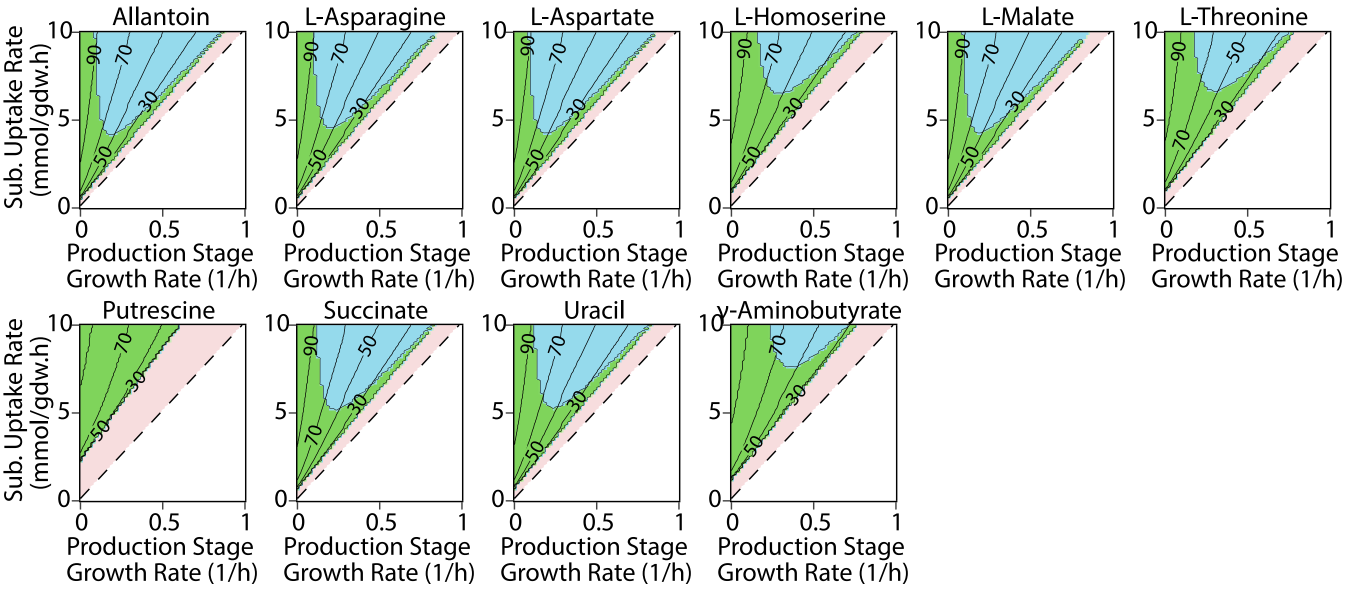


#### **e.** 5-Carbon Products

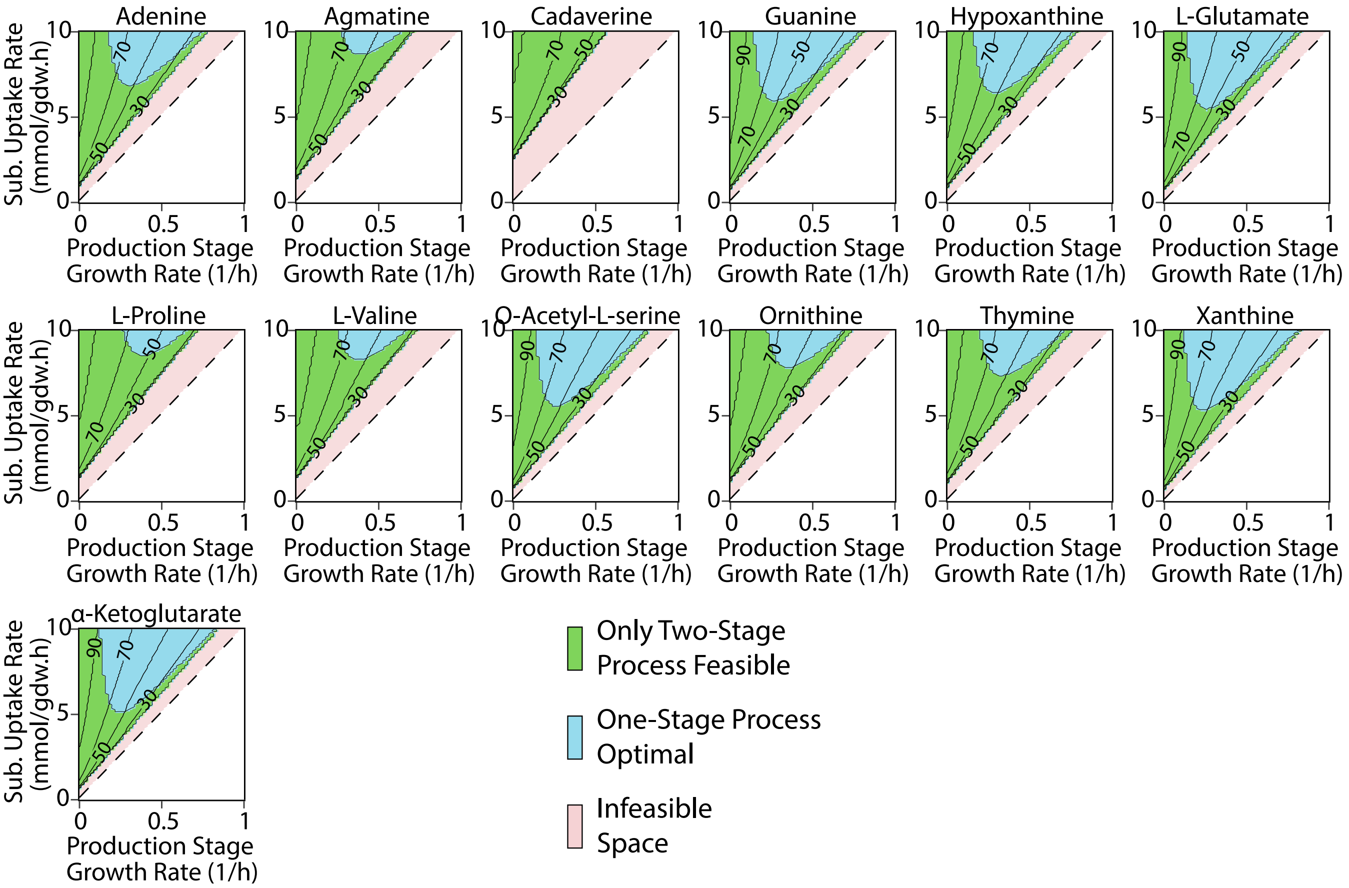


**Figure S9:** Comparison of TS and OS processes in *E. coli* at all possible substrate uptake rates during the production stage with fermentation objective III - maximize yield with a minimum productivity of 2 g/L.h for: **d.** 4 - carbon products. **e.** 5 - carbon products. The yields at each point within the feasible regions for TS and OS processes are represented by the contour lines as a % of the maximum yield

#### f. 6-Carbon Products

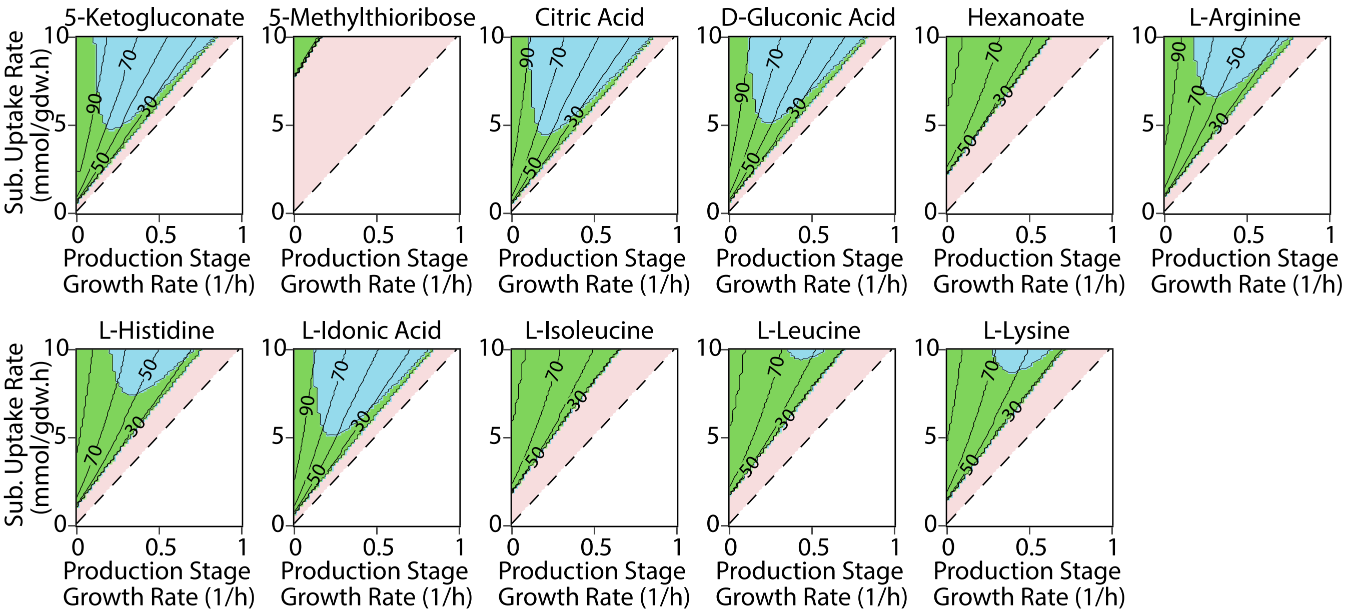


#### g. Products with > 6 Carbons

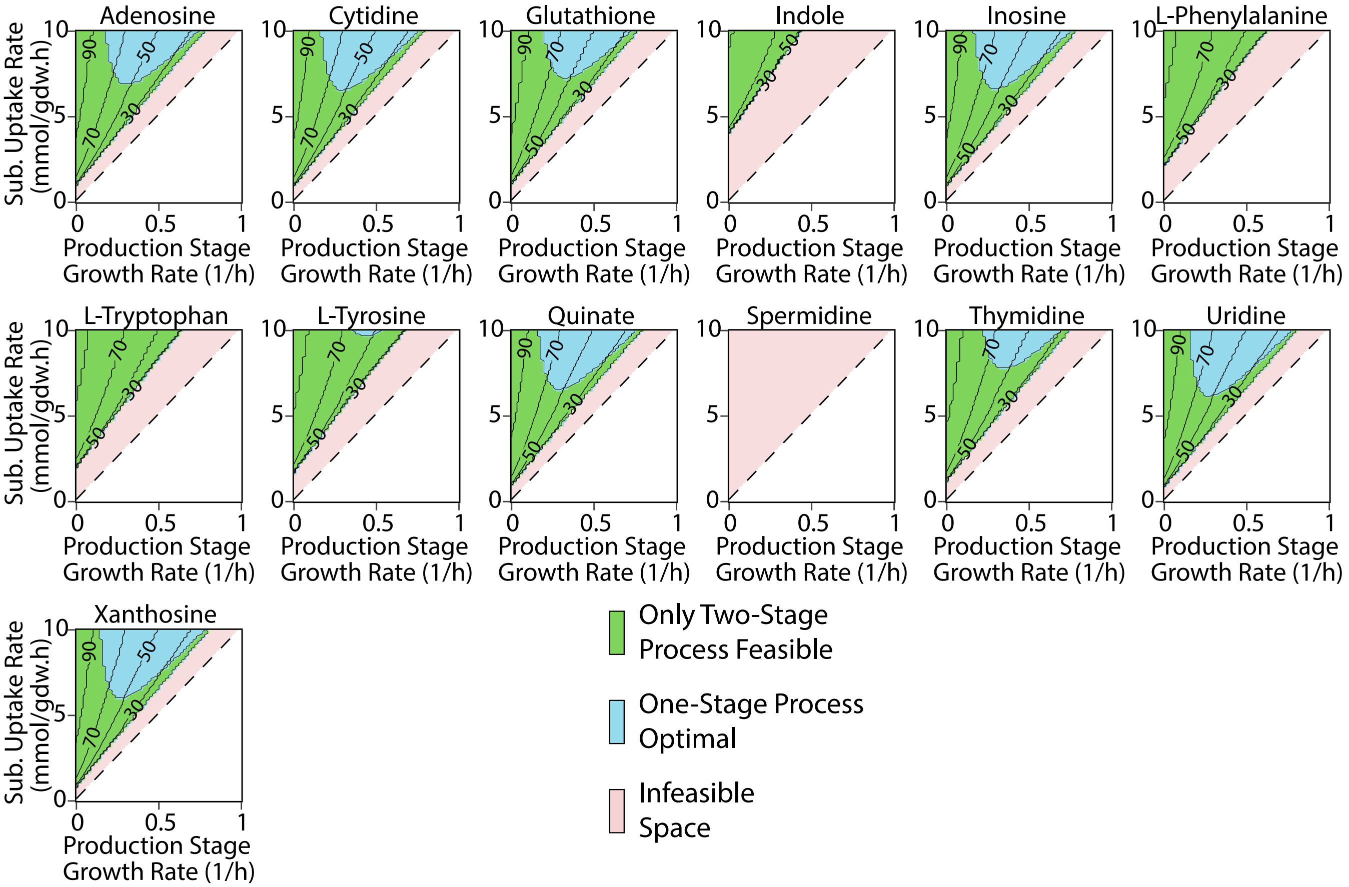


**Figure S9:** Comparison of TS and OS processes in *E. coli* at all possible substrate uptake rates during the production stage with fermentation objective III - maximize yield with a minimum productivity of 2 g/L.h for: **f.** 6 - carbon products. **g.** products with more than 6 carbon atoms. The yields at each point within the feasible regions for TS and OS processes are represented by the contour lines as a % of the maximum yield

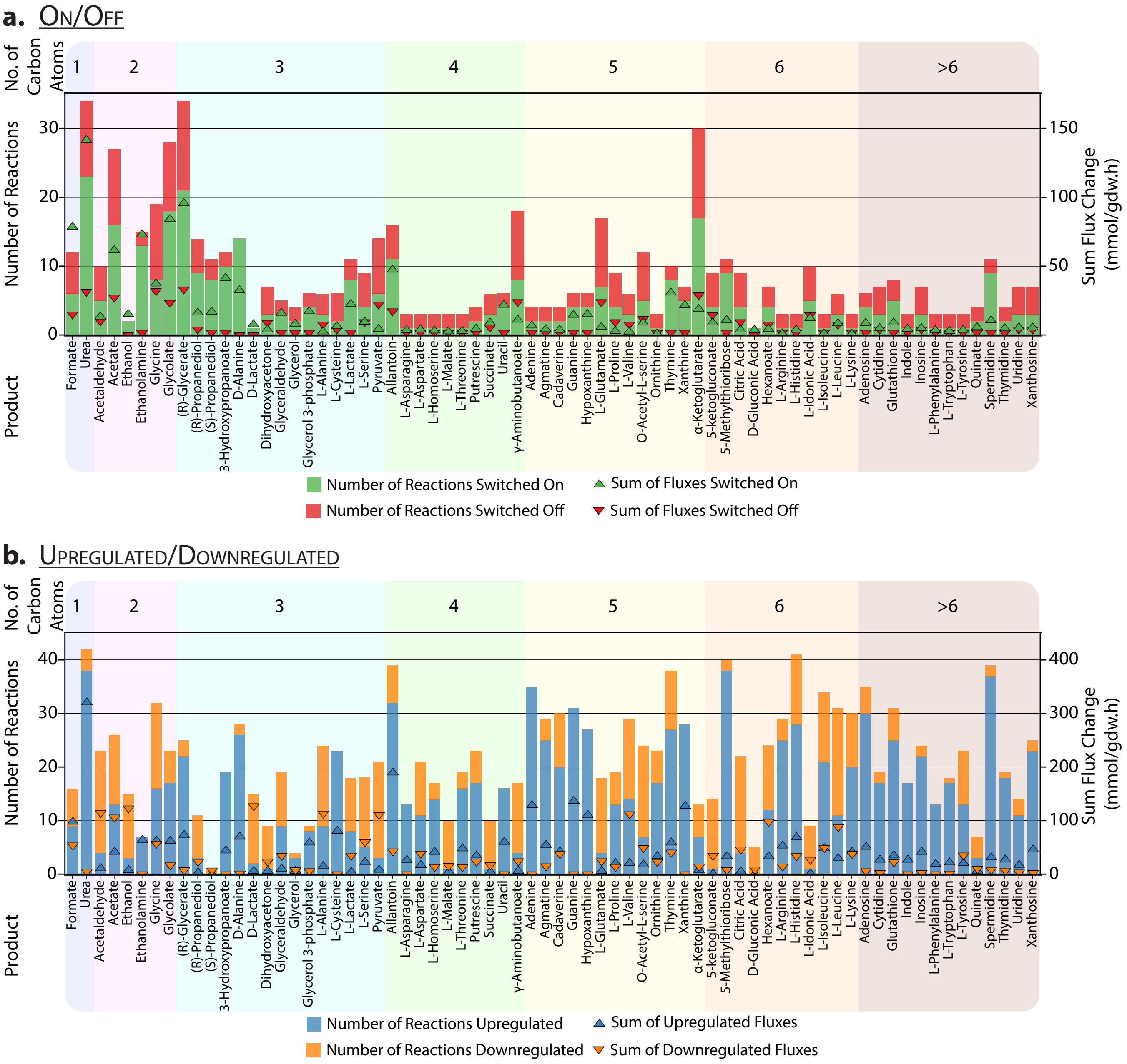


**Figure S10:** Number and sum of flux changes for reactions - **a.** switched on/off, **b.** upregulated/downregulated in two-stage strategies for native exchange metabolites in *E. coli* with Objective I - maximize productivity

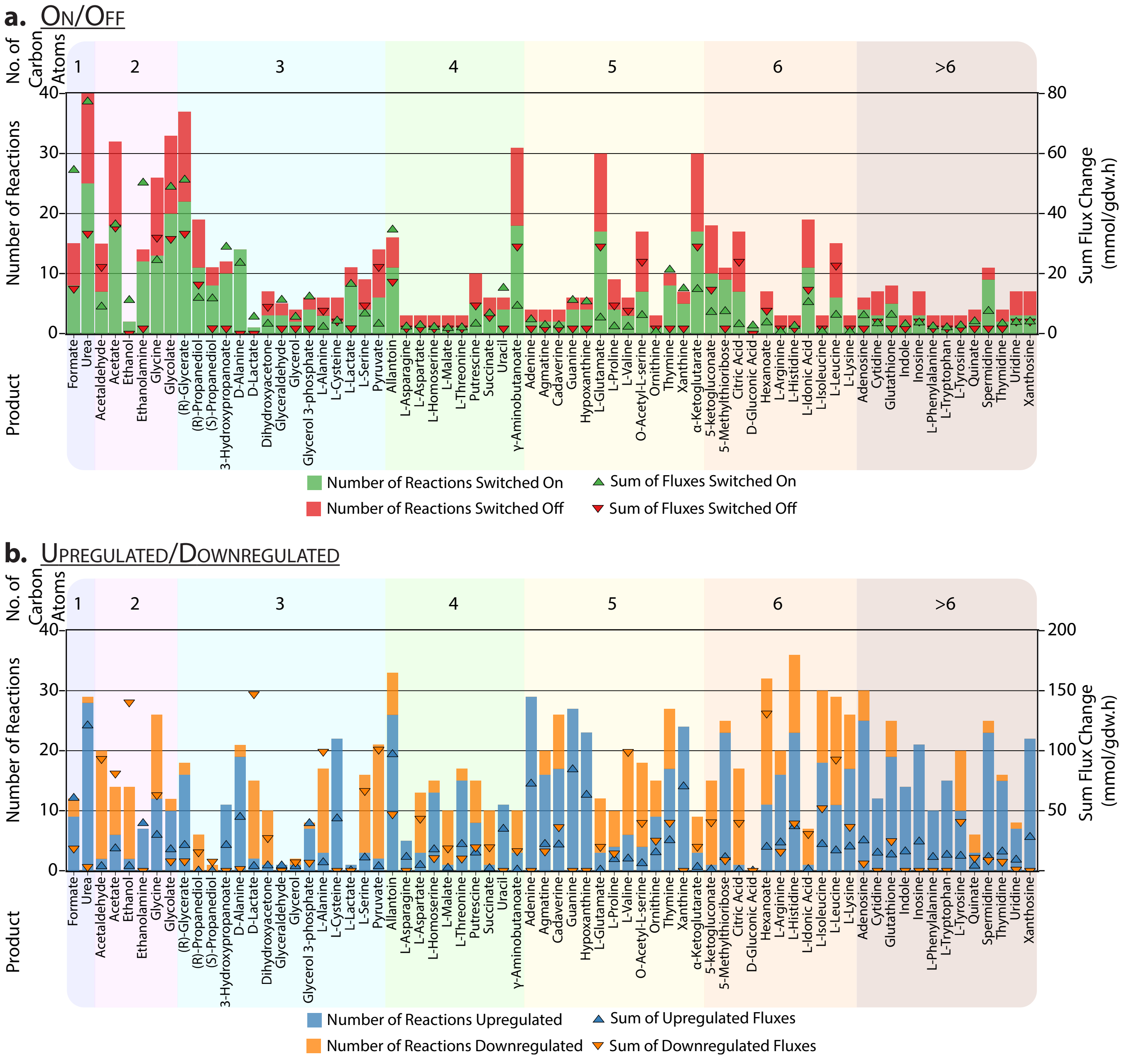


**Figure S11:** Number and sum of flux changes for reactions - **a.** switched on/off, **b.** upregulated/downregulated in two-stage strategies for native exchange metabolites in *E. coli* with Objective II - maximize productivity with a minimum yield of 75% of the maximum value.

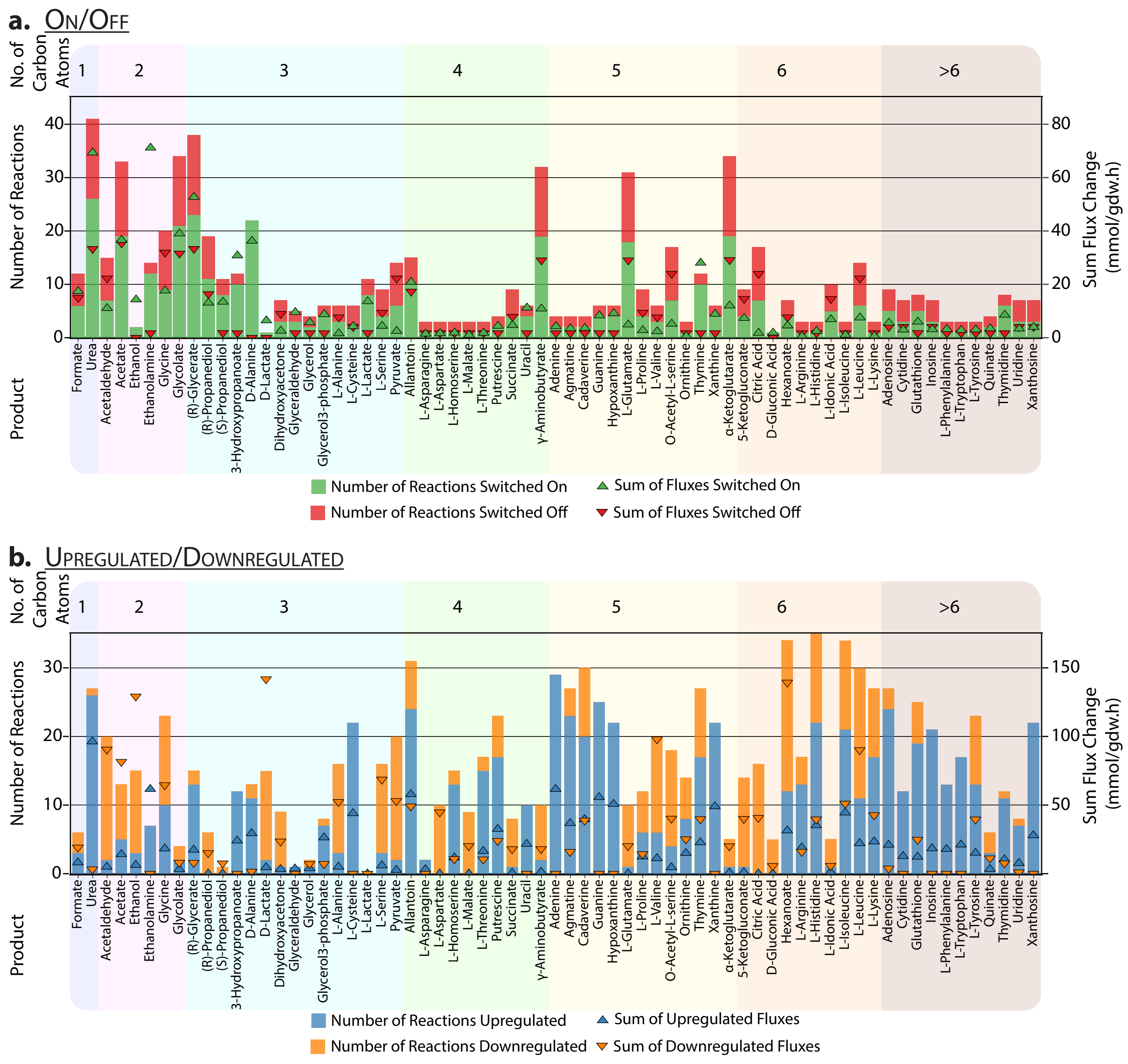


**Figure S12:** Number and sum of flux changes for reactions - **a.** switched on/off, **b.** upregulated/downregulated in two-stage strategies for native exchange metabolites in *E. coli* with Objective III - maximize yield with a minimum productivity of 2 g/L.h.

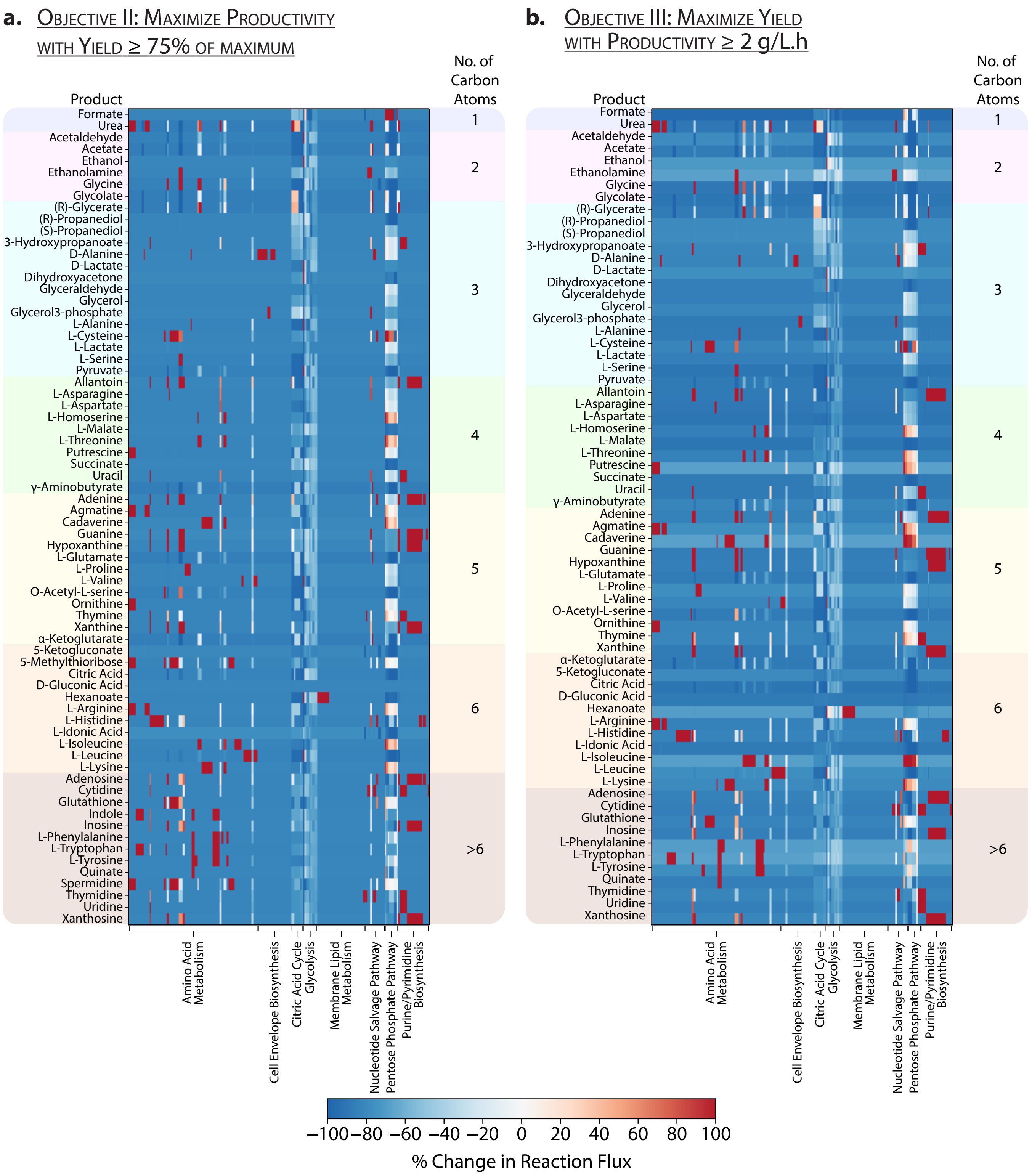


**Figure S13:** Percent change in reaction fluxes compared to wild-type fluxes during the production stage for native exchange metabolites in *E. coli* with: **a.** Objective II - maximize productivity with a minimum yield of 75% of the maximum value. **b.** Objective III - maximize yield with a minimum productivity of 2 g/L.h.

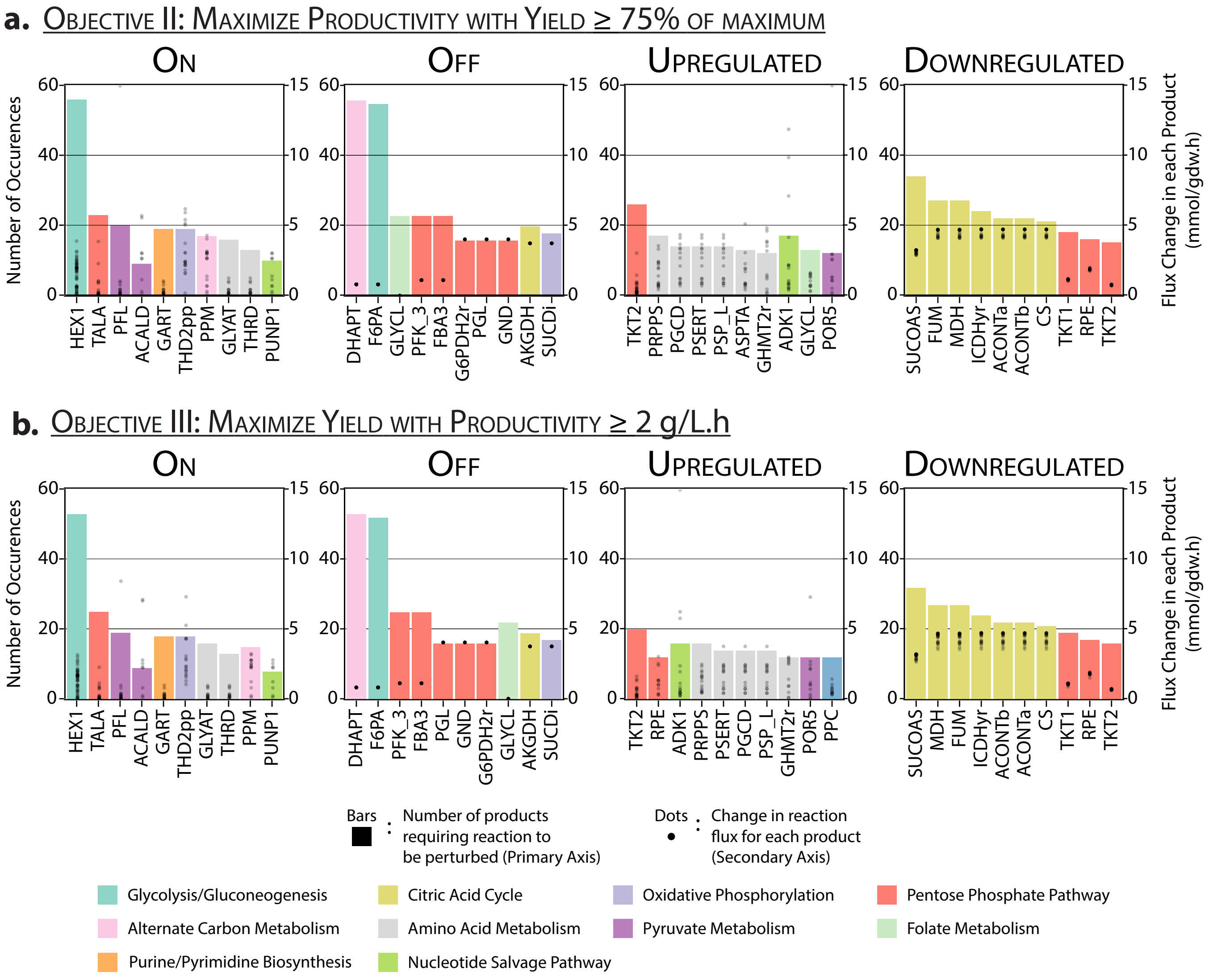


**Figure S14:** Reactions whose fluxes are changed for the most number of products in *E. coli* and their corresponding changes in flux values with: **a.** Objective II - maximize productivity with a minimum yield of 75% of the maximum value **b.** Objective III - maximize yield with a minimum productivity of 2 g/L.h, classified based on the type of change. Refer to Table S1 for reaction names and formulae
